# Single-cell metabolome and RNA-seq multiplexing on single plant cells

**DOI:** 10.1101/2025.05.20.655036

**Authors:** Moonyoung Kang, Anh Hai Vu, Abbie L. Casper, Rinho Kim, Jens Wurlitzer, Sarah Heinicke, Assa Yeroslaviz, Lorenzo Caputi, Sarah E. O’Connor

## Abstract

Plants produce valuable natural products used for a wide variety of industrial applications. Since these molecules have important applications in a variety of industrial sectors, there is enormous interest in elucidating the biosynthetic pathways that are responsible for the production of these compounds. Identification of the genes that comprise these biosynthetic pathways has been enabled by gene-to-metabolite networks that are generated from transcriptomic and metabolomic datasets. Recent advances in both single-cell RNA-seq (scRNA-seq) and single-cell mass spectrometry metabolomics (scMS) have enabled the measurement of either gene expression or metabolite levels in individual cells. However, these individual datasets can only be used to indirectly correlate gene expression levels with metabolite concentrations at the single cell level. Here we demonstrate that both scRNA-seq and scMS can be applied to the same plant cell, thereby enabling direct comparisons between gene expression and metabolite levels. This multiplexing approach reveals both qualitative and quantitative correlations between metabolite levels and biosynthetic gene expression in individual cells. This integrated approach sheds light on the underlying processes driving complex plant biosynthesis.

## Introduction

Plants produce an extraordinary array of complex natural products. Since these molecules have important applications in pharmaceutical, agricultural and nutritional sectors, there is enormous interest in elucidating the biosynthetic pathways that are responsible for the production of these plant-derived compounds. Identification of the genes that comprise these biosynthetic pathways has been greatly facilitated by the availability of transcriptomic and metabolomic datasets. These data are used to construct gene-to-metabolite networks that allow correlation of gene expression with metabolite levels. These correlation analyses have led to the discovery of many biosynthetic genes, transporters and regulatory elements from plant pathways (1–6).

Single cell transcriptomic (scRNA-seq) approaches have enabled a step-change in the resolution to which gene expression profiles can be measured (7, 8). scRNA-seq is now well-adapted for a number of plant systems, and, since plant biosynthetic genes of specialized metabolism are typically localized to only a few specific cell types, scRNA-seq is particularly well-suited for analysis of natural product pathways (9–11). However, generation of accurate gene-to-metabolite correlations at the single cell level requires, in addition to scRNA-seq datasets, corresponding single cell mass spectrometry (scMS) data. We recently reported a scMS method for analysis of single plant protoplasts and subsequently demonstrated how the resulting scMS data could be interpreted alongside scRNA-seq to facilitate the discovery of biosynthetic genes (10, 12). However, correlations between these two distinct datasets could only be made indirectly as the generation of gene-to-metabolite networks using these data was not possible.

We envisioned that rigorous gene-to-metabolite correlations could be achieved through a multiplexed approach in which both scMS and scRNA-seq data are obtained from the same individual plant cell. Here, we report a method by which a single plant protoplast can be analyzed by RNA-seq and metabolomics. Using *Catharanthus roseus*, a medicinal plant known for producing an array of terpenes, alkaloids and phenylpropanoid natural products as a proof-of-concept (13), we demonstrate qualitative and quantitative correlations between metabolite levels and biosynthetic gene expression at the single-cell level. These datasets show how the intermediates and end-products in the pathway of the alkaloid anhydrovinblastine, a precursor to the anti-cancer agent vinblastine, are located relative to the biosynthetic genes. Overall, this integrated method offers a powerful tool for understanding the spatial organization of metabolite biosynthesis in plants and may accelerate the discovery of genes involved in specialized biosynthetic pathways.

## Results

### scRNA-seq - scMS multiplexing workflow and data acquisition

A recently developed scMS method for profiling metabolites in plant single cells relies on sorting plant protoplasts on a SIEVEWELL™ chip (10, 12). After being trapped into an individual well of the chip, each protoplast is picked using a robot system (CellCelector™ Flex), and transferred into a small volume of water in a microtiter plate. The resulting osmotic shock causes immediate lysis. After addition of solvent and internal standard, an aliquot of this solution is analyzed using a standard UPLC-MS instrument. Because this method includes a chromatographic step, metabolites can be structurally identified by comparison of retention time and MS/MS fragmentation with authentic standards. Additionally, using calibration curves for these standards, along with cell diameter measurements obtained from images captured during the picking process, metabolite concentrations can be calculated for each cell. We reasoned that this scMS workflow could be adapted for multiplexing with a plate-based scRNA-seq approach (14). To preserve RNA integrity, protoplasts were dispensed into 8 μL of RNase-free water containing RNase-inhibitor. Following lysis, each well’s contents were split: half was used to generate a replica plate for RNA-seq, stored at –80°C prior to cDNA library preparation using the SMART-seq2® plate-based protocol, while the other half was used for UPLC-MS analysis (Fig. 1A).

**Figure 1.**
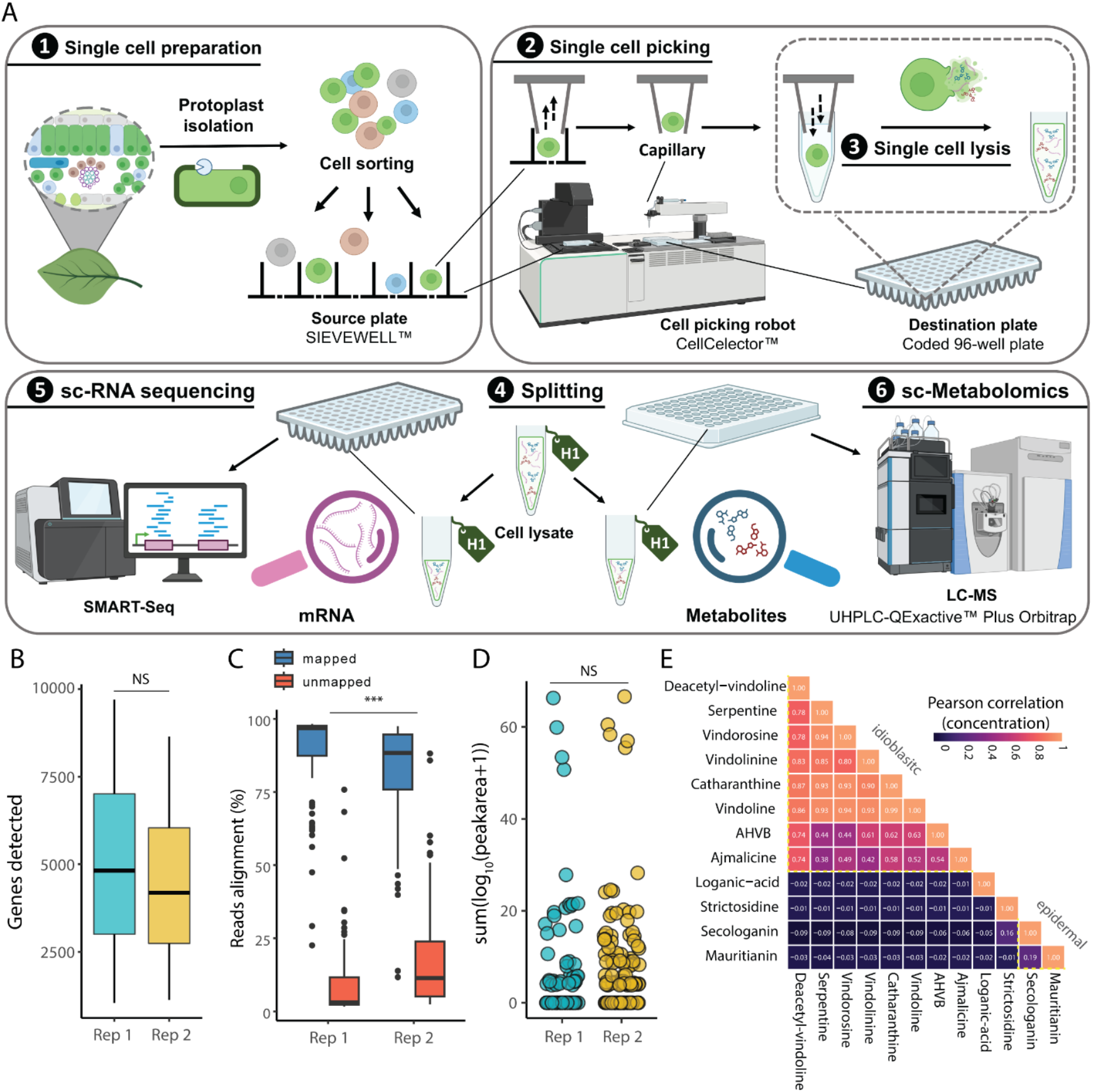
Multiplexing of single plant protoplasts. **A.** Workflow for simultaneous single cell transcriptomics and metabolomics of plant protoplasts. **B.** Number of genes detected in single cells. **C.** Sequencing read alignment ratio to genome. **D.** Number of metabolites detected in single cells. Statistical differences between the two groups were calculated by performing two-sided Kolmogorov-Smirnov (KS) tests (***, *P*<0.01; NS, not significant). For boxplots, the box represents the interquartile range (IQR), and the whiskers indicate highest and lowest points within 1.5× IQR. **E.** Summary of metabolomics profile, performed as described in a previous study (12). Pearson correlation coefficients between concentrations of 12 analytes are visualized based on color scale.

Protoplasts were isolated from young leaves of the medicinal plant *Catharanthus roseus*. To ensure sufficient number of cells for the experiment, a total of four 96-well plates were collected on separate days. Two plates were prepared each time from freshly isolated protoplasts derived from leaves of two individual plants. Image analysis from the cell picking process confirmed that 289 cells were successfully picked, while 12 wells contained cell doublets and 95 wells were empty. The lysates from these 289 cells were subjected to both transcriptomic and metabolomic analysis. Following initial quality assessment of the scRNA-seq data, cells with fewer than 1000 detected genes were excluded from the dataset, resulting in a total of 193 (66%) retained for further analysis. Across the entire dataset, a total of 21,258 genes were identified, with each cell expressing an average of 5000 genes (Fig. 1B) with a mapping rate greater than 80% (Fig. 1C). No significant batch variation was observed when we combined the data from the cells collected on the two different days (Fig. S1).

**Scheme 1.**
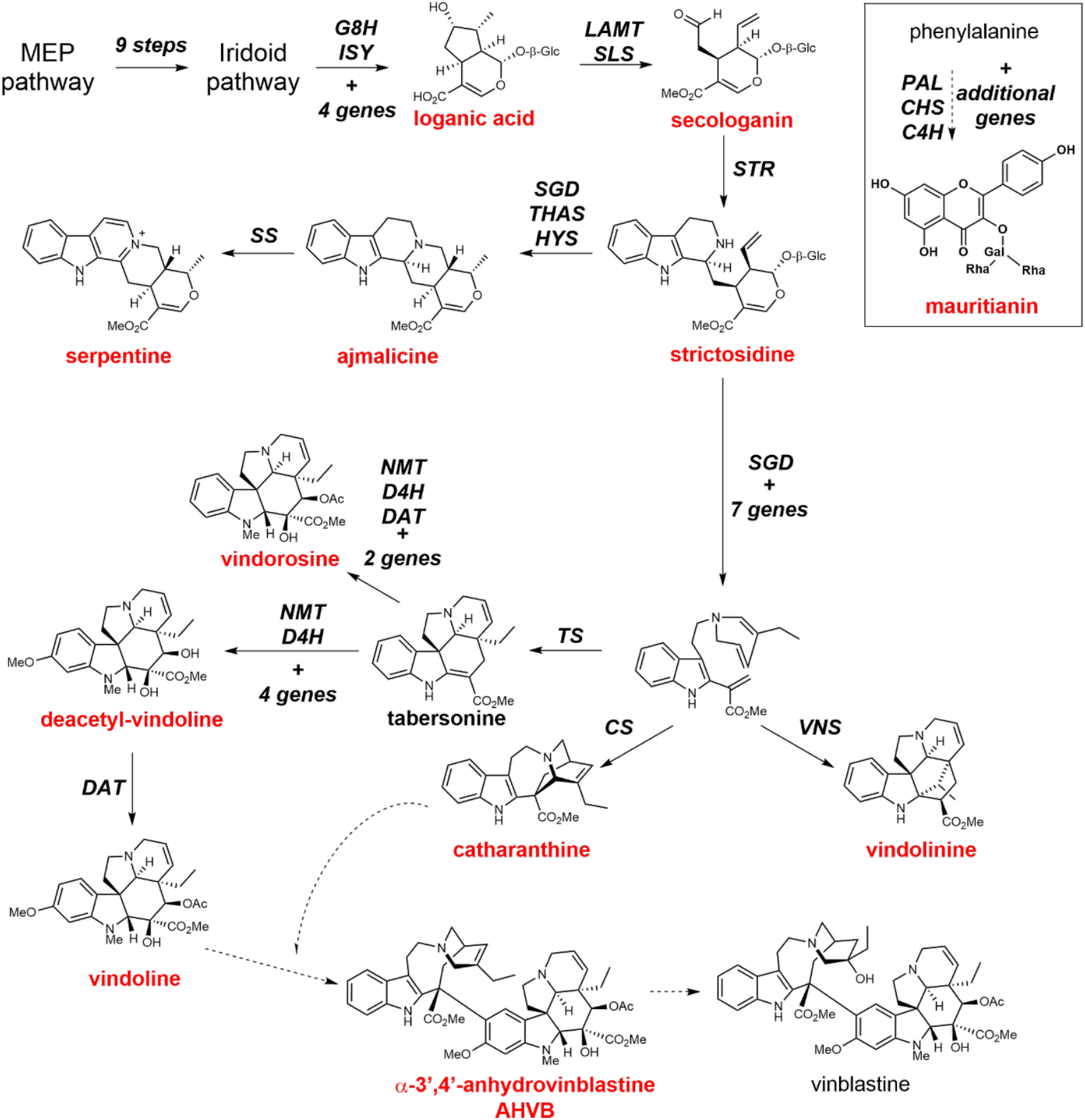
Pathway to metabolites of *Catharanthus roseus.* Key metabolites targeted in this study are highlighted in red text. Selected biosynthetic genes are indicated in black. Dashed lines indicate that biosynthetic genes for these steps are unconfirmed. intermediates are shown. Mauritianin, which is biosynthetically unrelated to the other metabolites targeted, is shown in the inset box. A complete biosynthetic pathway including the full name of the enzymes is shown in Supplementary Scheme 1.

We next analyzed the scMS data from the same set of 193 single cells. Targeted mass spectrometry was employed to quantify 14 metabolites of interest, for which authentic standards are available. Under these conditions, intracellular concentrations were successfully quantified for 12 of the 14 metabolites (Table S1). Notably, several key intermediates in the anhydrovinblastine biosynthetic pathway, including the iridoid loganic acid, the seco-iridoid secologanin and the alkaloids catharanthine and vindoline, were detected at high concentrations, reaching millimolar levels in some cells (Scheme 1). Anhydrovinblastine itself accumulated in micromolar concentrations, while vinblastine, typically present in much lower levels, was not observed in this limited sample set. Additional intermediates in the pathway, such as strictosidine and deacetylvindoline, were observed at low micromolar concentrations. We also quantified several related alkaloids–vindorosine, vindolinine, ajmalicine, and serpentine–which are biosynthetically related to, but not direct intermediates of, anhydrovinblastine (Scheme 1). In addition to these alkaloids, we quantified the flavonoid mauritianin, a kaempferol diglycoside, which is one of the major flavonoids found in *C. roseus* leaf. All metabolites were quantified across the 193 cells analyzed (Figs. S2, S3). Comparison of data from the two separate collection days showed strong consistency between the biological replicates and was in agreement with previously reported scMS results for this species (Table S2).

Correlation analysis of the scMS dataset revealed that all nitrogen-containing alkaloids, with the exception of strictosidine, consistently co-occurred within the same cells. Based on the presence of vindoline, these cells were tentatively identified as idioblast cells (10, 12, 15) (Fig. 1E). Strictosidine is rarely detected in leaf protoplasts, but when present, it co-occurs with its precursor secologanin (Pearson correlation r >0). Secologanin is predicted to localize primarily in the epidermal cell. Although flavonoids in *C. roseus* are also predicted to be localized to epidermal cells (16), mauritianin was detected only in a subset of cells that also accumulated secologanin (12). Loganic acid, which is synthesized in internal phloem-associated parenchyma (IPAP) cells, was used as a marker to tentatively classify loganic acid-accumulating cells as IPAP cell types.

### Cell clustering based on gene expression and metabolite accumulation

A UMAP projection was generated from the scRNA-seq data (RNA-UMAP) and cell types were annotated based on previously validated marker genes (17–20). The gene *NLTP2* was used as a marker gene for epidermis, while *CB21*, known not to be expressed in epidermal cells, was used to validate this assignment (18) (Fig. S4A). *DAT* and *D4H* transcripts served as markers for idioblasts. Cells were annotated as IPAP when either *G8H* or *ISY* transcripts, which are reported to be highly specific to this cell type were detected (19, 20). In the RNA-UMAP, cells expressing epidermal and idioblast markers formed well-defined clusters (Fig. S4A). In contrast, no distinct cluster corresponding to IPAP cells was observed. *ISY* was expressed at low levels in only two cells, while *G8H* was detected in just six cells, all located outside the idioblast and epidermal clusters (Fig. S5). We assigned these cells as IPAP type, even though *G8H* and *ISY* were expressed at low levels. However, in actuality, this sample of protoplasts may not have included IPAP cells, which are known to be a rare cell type, comprising ca. < 4 % of the total leaf cell population, based on previous studies (21).

We next projected cell type-specific metabolites onto this RNA-UMAP (Fig. S6A). As expected, secologanin localized to the same region as the epidermal marker, consistent with its predicted epidermal origin. Similarly, the flavonoid mauritianin also mapped to the epidermal region. In contrast, alkaloids, e.g. serpentine and anhydrovinblastine, correlated to the idioblast region of the RNA-UMAP, in line with the known localization of these compounds. Loganic acid, which is predicted to be synthesized in IPAP cells, showed no correlation with cells expressing IPAP marker genes. This is consistent with our earlier observation that a distinct IPAP cell cluster was not evident on the RNA-UMAP. Given that loganic acid must be transported from IPAP cells to epidermal cells (22, 23), where it is converted to secologanin, we initially hypothesized that its presence might coincide with epidermal cell markers. However, loganic acid did not correlate with the epidermal cell either. Instead, the seven cells in which loganic acid was detected were primarily located in the parenchyma cell region of the RNA-UMAP. It is possible that the high concentrations of loganic acid observed (average intracellular concentration of ca. 40 mM) represent molecules that have been exported from the IPAP cells and stored in a distinct, yet uncharacterized, cell type. The cell type(s) in which loganic acid accumulates could not be confidently annotated due to the limited availability of specific gene markers for plant cell types.

To further investigate metabolite distribution across cell types, we generated a UMAP based on the 12 quantified metabolites rather than gene expression data (MET-UMAP, Fig. 1E). This analysis revealed distinct clusters of cells enriched in specific compounds, including alkaloids (e.g. serpentine, vindoline, anhydrovinblastine), secologanin, mauritianin or loganic acid (Figs. 2B, S4B). We then compared the RNA-UMAP and MET-UMAP to assess the spatial correlation of selected gene-metabolite pairs (Fig. 2C). Notably, 8 out of 9 (89 %) cells that accumulate serpentine also expressed the idioblast cell markers *D4H* and *DAT*. Similarly, secologanin showed strong correlation with the epidermal marker *NLTP2*, with 29 out of 38 (76 %) cells that accumulate secologanin also expressing this marker (Fig. 2D, Table S3). Remarkably, the MET-UMAP revealed two distinct clusters within the epidermal population: one accumulating secologanin alone, and the other accumulating both secologanin and the flavonoid mauritianin. Cells in both clusters corresponded to *NLPT2*-expressing cells in the RNA-UMAP, suggesting the existence of two metabolically distinct cell subtypes. Transcriptomic comparison of these two epidermal subpopulations revealed several differentially expressed genes, including upregulation of *KCS* and lipid transfer proteins in the mauritianin-accumulating cells. However, expression levels of predicted flavonoid biosynthetic genes (e.g. *PAL, CHS, C4H*) did not differ significantly between the two groups (Figs. S10, S11), suggesting that the genetic basis underlying their metabolic differentiation may be more complex than anticipated.

**Figure 2.**
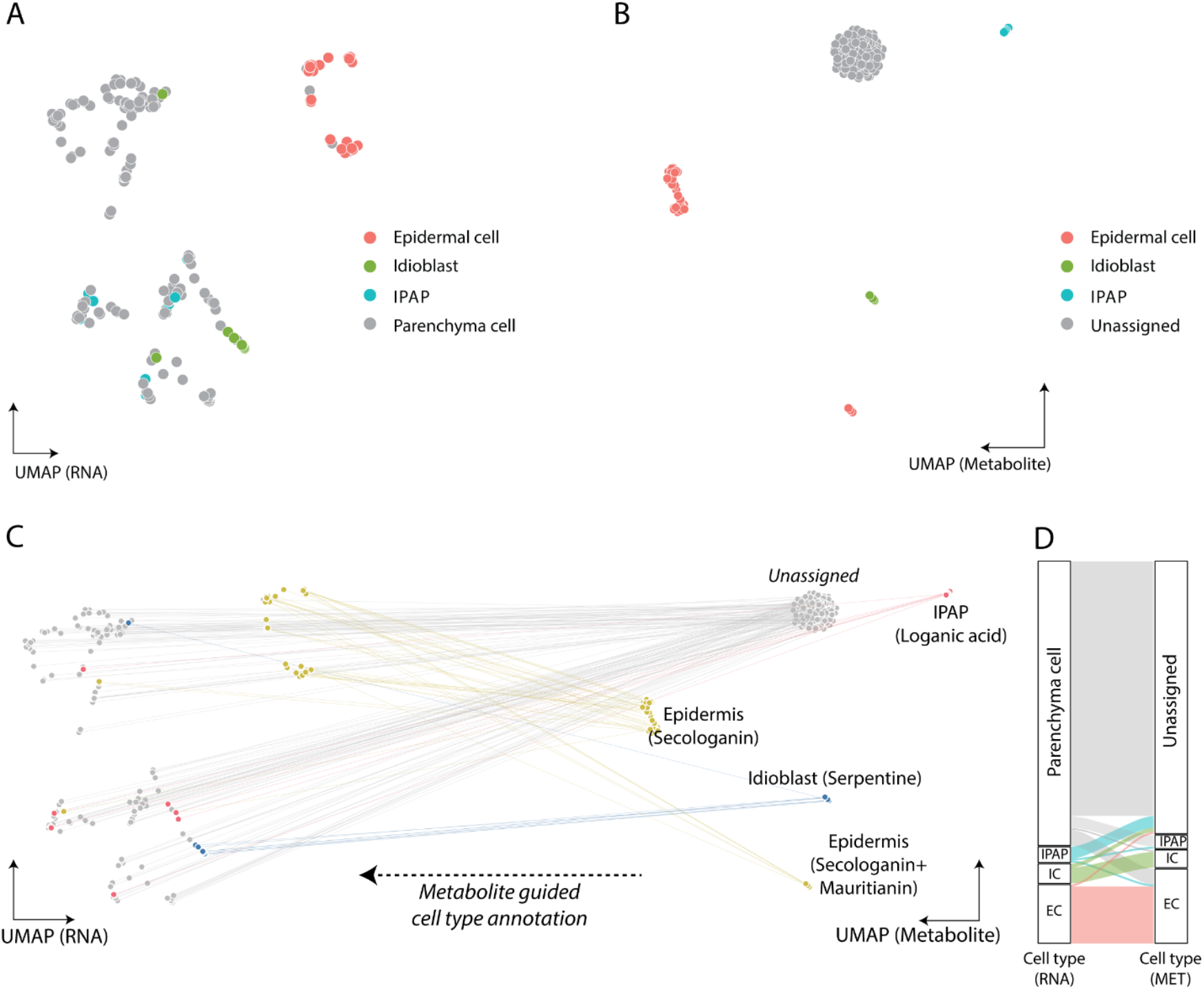
UMAP showing clustering of cell types by gene expression and metabolite expression.**A.** Marker gene guided cell type annotation in RNA-UMAP plot. **B.** Targeted analyte guided cell type annotation in MET-UMAP plot. **C.** These two different UMAP plots were connected based on their corresponding well identities, where cells are colored with annotations based on representative metabolite accumulation (serpentine, secologanin, mauritianin, loganic acid). **D.** Sankey plot showing similarities and differences when cells are annotated with gene markers or metabolites. The height of the plot is proportional to the number of cells. The left side represents the marker gene-based cell type annotation, and right side represents metabolite-based annotation.

### Quantification of gene-metabolite correlations

Since both gene expression levels and selected metabolite concentrations were quantified in this study, we were able to assess gene-metabolite relationships using Spearman correlation analysis (Fig. 3A). Specifically, we calculated the correlation between the occurrence of specific metabolites and the expression levels of all detected genes across the dataset. For several metabolites we observed strong correlations with the expression of known biosynthetic genes. For example, vindoline showed high correlation with the expression of the genes *D4H, DAT* and *NMT*–the final genes in its biosynthetic pathway (Scheme 1). This strong correlation is consistent with the fact that vindoline is stored in the same cell type in which it is synthesized. Similarly, anhydrovinblastine, a downstream, derivatized product of vindoline, that is also localized in the idioblasts, showed strong correlation with *D4H, DAT* and other idioblast-enriched genes.

**Figure 3.**
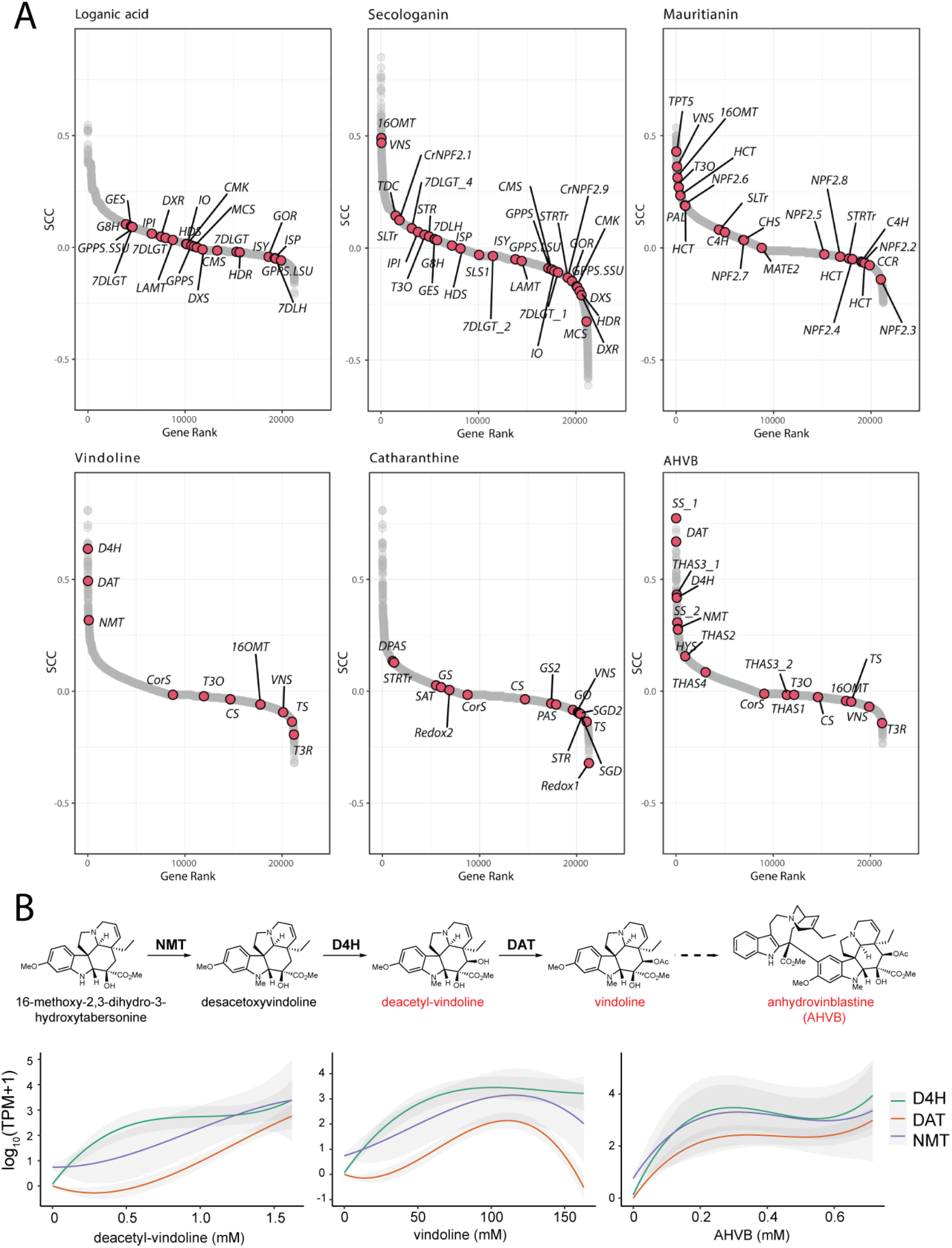
Quantitative gene to metabolite correlation.**A.** Spearman correlations of all genes were plotted according to their correlation rank with each of the listed compounds. **B.** Trend lines (local regression with 95% confidence interval) of NMT, D4H, DAT expression levels (TPM) are plotted according to ascending concentration of each listed compound.

In contrast, some biosynthetic genes showed weak or no correlation with alkaloid accumulation, likely due to extensive transport between cell types. While alkaloids are ultimately stored in idioblasts (Fig. S8), many upstream steps in their biosynthetic pathway take place in epidermal cells (Figs. S7C, S9). For instance, *T3O* and *T3R*, which function early in vindoline biosynthesis and are localized to the epidermis, did not correlated with vindoline presence (Fig. 3A). Similarly, catharanthine is synthesized in epidermal cells but subsequently transported to idioblasts for storage. As a result, we observed no meaningful correlation between catharanthine and the expression of its biosynthetic genes (e.g. *DPAS, PAS, CS)* (Fig. 3A).

We also examined correlations for secologanin and mauritianin, two metabolites that are expected to be both synthesized and retained within epidermal cells. While the correlation with specific biosynthetic genes was modest (e.g. *LAMT* for secologanin, *C4H* for mauritianin), both metabolites showed strong correlation with a range of genes known to be expressed in epidermal cells, such as *16OMT* and *VNS*. Thus, although direct correlations with individual biosynthetic genes were limited, these metabolites aligned well with markers for the correct cell type. In contrast, the iridoid loganic acid, which is synthesized in IPAP cells and then transported to epidermal cells for conversion into secologanin, did not show significant correlation with either IPAP- or epidermis-specific genes, as discussed above (Figs. 2D, 3A). This reinforces the hypothesis that the large pools of loganic acid observed by scMS likely reflect transport to a third, unidentified cell type where it is stored or transported through.

We next examined the relationship between absolute metabolite concentrations and the expression levels of specific biosynthetic genes. As a proof-of-concept, we focused on the idioblast localized metabolites deacetylvindoline, vindoline and anhydrovinblastine, and their corresponding biosynthetic genes *NMT, D4H* and *DAT*, as these gene-metabolite pairs showed strong qualitative correlations. We observed that the expression of these biosynthetic genes generally increased in parallel with rising alkaloid concentrations, reaching peak expression at approximately 100 mM metabolite level (Fig. 3B). Beyond this point, gene expression levels began to decline slightly. We speculate that this pattern may reflect a developmental trajectory of idioblasts, transitioning from active biosynthesis to storage phase, in which mature cells accumulate high alkaloid concentrations while exhibiting reduced gene expression. Alternatively, these high levels of alkaloid may lead to feedback inhibition of gene transcription. However, the relatively small number idioblasts captured in this dataset limits our ability to test these hypotheses with statistical confidence.

### Relationship of transporters and metabolites

Given the extensive intercellular transport involved in these metabolic pathways, we also assessed the correlation between metabolites and expression of the genes encoding previously identified transporters. NPF2.4, 2.5, 2.6 were shown to transport loganic acid, loganin and secologanin in an *in vitro* assay (24). Our data suggest that NPF2.4 is the physiologically relevant transporter that imports iridoids into the epidermal cells, as its expression profile closely mirrors the accumulation of secologanin in this cell type (Fig. 4A). TPT2, which has been reported to export catharanthine from epidermal cells to cuticle layer (25), was not detected in any of the 193 cells analyzed, suggesting it may not play a significant role in catharanthine transport in planta. However, a homologous gene, TPT5, was expressed in a subset of epidermal cells, though its function in unknown (25). Although these are the only inter-cellular transporters characterized to date in *C. roseus*, the integrated transcriptomic and metabolomic dataset presented here provides a valuable resource for identifying additional candidate transporters involved in specialized metabolites trafficking.

**Figure 4.**
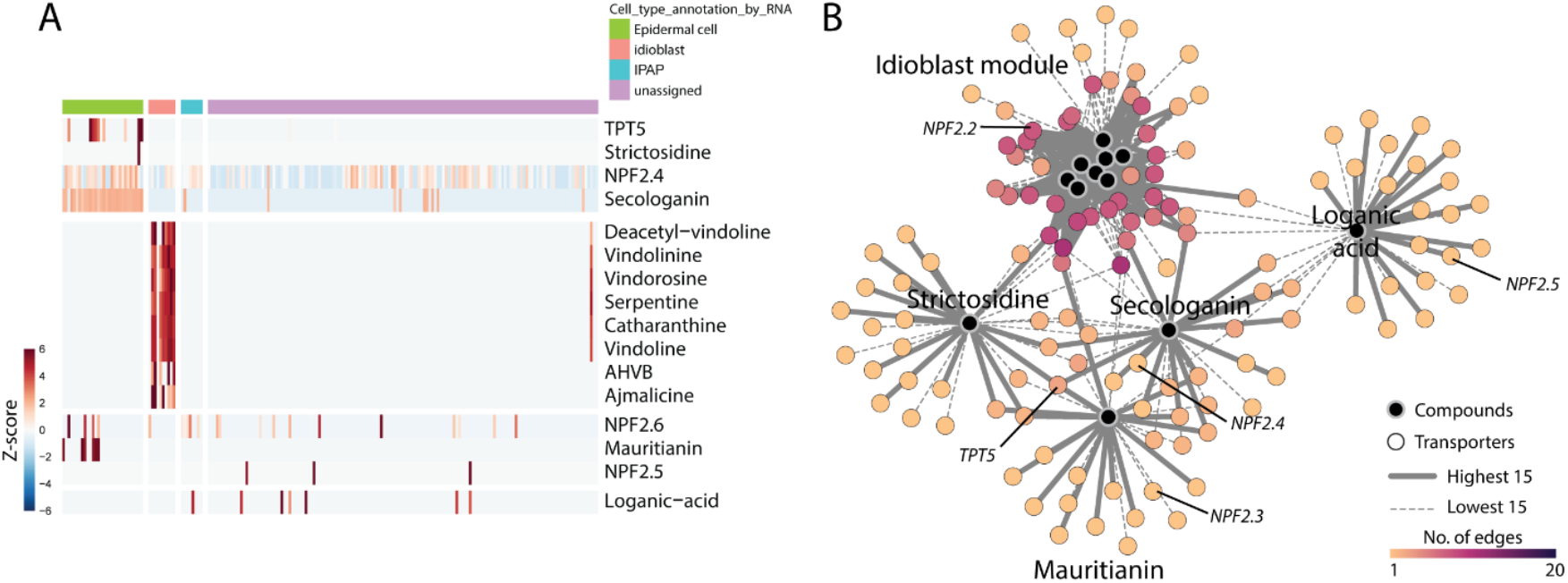
Transporters and metabolites.**A.** Heatmap showing coexistence of known intercellular transporter-metabolite pairs. Z-scores of each gene expression and metabolite accumulation in single cell was colored and hierarchically clustered. **B.** Expected network of metabolites and putative importers/exporters based on correlation. Transporters with the top 15 highest and bottom 15 lowest correlation values are linked to metabolites in the network. The number of connections for each transporter is represented by a color scale. Transporters are annotated if they were studied or mentioned in literatures.

## Discussion

Here, we demonstrate that small molecule mass spectrometry and RNA-seq can be simultaneously applied to a single plant protoplast. Multiplexing approaches that integrate metabolomics at single-cell level are not widely reported, due to the challenges associated with observing and annotating small molecules in single cells. To date, the only other reported single-cell multiplexing approach involving metabolomics employed a nano capillary to manually sample just fifteen mammalian cells (26). While that study represents a technically rigorous and elegant approach, the method presented here is specifically adapted for plant cells, offers significantly higher throughput, and enables accurate quantification of individual metabolites.

Both scMS and scRNA-seq provide valuable insights into cellular metabolism, each with its own limitations. Single cell metabolomics reveals the identity and quantities of selected molecules in cells, but in the absence of gene expression data, the biological interpretation of these measurements remains limited. Conversely, single cell RNA-seq data provides information about the location of biosynthetic genes, but cannot directly link genes to their corresponding metabolites. By simultaneously profiling the transcriptome and metabolome of a single cell, we are able to directly associate cell type markers and biosynthetic genes with the presence of specific metabolites. This integrated dataset clearly distinguishes between metabolites that are retained in the site of synthesis and those that are transported to a different cell type. For example, our data showed that the metabolic pools of loganic acid that we observe are not stored in either idioblast or epidermal cells, suggesting that this alkaloid pathway intermediate is transported to and stored in an unexpected third cell type. In contrast, metabolites such as the late-stage alkaloid intermediates, which are both synthesized and stored in the idioblasts, show strong quantitative correlations between gene expression and metabolite concentration. Importantly, this approach allows us to track gene expression dynamics along the concentration gradient of specific metabolites at single-cell resolution. Additionally, the use of a plate-based scRNA-seq method enables the detection of a greater number of genes per cell compared to droplet-based approaches, yielding a deep transcriptomic profile that is particularly suitable for gene mining.

It is important to note that this method is limited to cells that can be isolated as protoplasts. While previous studies have shown that natural product biosynthetic pathways remain functional or even upregulated after protoplast formation (10), certain biological processes are inevitably affected by the protoplasting procedure, and may not be accurately captured using this approach. Moreover, the relatively low throughput of this method (ca. 200 cells per experiment) means that rare cell types may be missed. Another limitation lies in the sensitivity of the mass spectrometry instrumentation. Only metabolites that accumulate to levels above the limit of detection of the method can be measured whilst short-lived biosynthetic intermediates are often not detected. Despite these constraints, this approach provides a powerful mean to correlate gene expression with metabolite localization at single-cell resolution, thereby adding an important dimension to our understanding of plant metabolism. We envision that these datasets will aid in the identification of biosynthetic genes involved in plant metabolism. Furthermore, the application of this approach to other plant systems will enable to compare and contrast how gene expression and metabolite storage is conserved across plant metabolic pathways.

## Materials and methods

### Plant growth conditions

*Catharanthus roseus* (*C. roseus*) plants (Sunstorm Apricot cultivar) were germinated and grown in a York chamber at 23 °C, under a 16 h:8 h, light: dark cycle. Leaves of *C. roseus* were harvested from approx. 2-month-old plants just before the experiment.

### Chemicals and reagents

Commercially available analytical standards used in this study included: loganic acid (Extrasynthese), secologanin (Sigma Aldrich), mauritianin (BIOMOL GmbH), ajmalicine (Sigma Aldrich), catharanthine (Abcam), deacetylvindoline (Toronto Research Chemicals Inc), anhydrovinblastine disulfate (Toronto Research Chemicals Inc.), vinblastine sulfate (Thermo Scientific Chemicals), serpentine hydrogen tartrate (from Sequoia Research Products Ltd.), ajmaline (Extrasynthese), tabersonine (TCI, Tokyo Chemical Industry Co. Ltd.), vindolinine (Advanced ChemBlocks Inc.), vindoline (Acros Organics). Strictosidine and vindorosine were synthesized and isolated in our laboratory, as previously reported (12, 27).

For protoplast extraction, cellulase Onozuka R-10 and macerozyme R-10 were obtained from SERVA, while pectinase, mannitol, KCl, MES, and Bovine serum albumin were purchased from Sigma Aldrich. All solvents used in this study were of UHPLC/MS grade.

### Single cell collection

#### Protoplast isolation

Young leaves (2~3 cm) of *C. roseus* plant were rinsed with distilled water, cut in strips with a surgical blade and immediately submerged in digestion medium (2 % (w/v) Cellulase Onozuka R-10, 0.3 % (w/v) macerozyme R-10, 0.1 % (w/v) pectinase, and 0.1 % (w/v) bovine serum albumin dissolved in Mannitol-MES (MM) buffer). MM buffer contained 0.45 M mannitol and 20 mM MES, pH 5.7-5.8. Digestion medium was infiltrated into the leaf strips by applying vacuum for 15 min (200 mBar), and protoplasts were released from the leaf strips while shaking for 2.5 h at room temperature. The protoplast suspension was strained through 40 μm sieves to remove large debris, and transferred to 15 mL round bottom tubes. Protoplasts were centrifuged at 70 *g* for 5 min and washed with corresponding MM buffer three times.

#### Single cell picking

After washing, the protoplasts were resuspended in a small volume of MM buffer. After FDA staining, their concentration and cell viability were checked under microscope. Single cell picking was performed as described previously (12). Briefly, approximately 10,000 protoplasts were dispensed onto a Sievewell™ (Sartorius) with cell-size micropores (50 μm) to capture single cells by gentle suction-induced sedimentation. The CellCelector™ Flex (Sartorius) cell picking robot was used to transfer the targeted cells from the Sievewell™ to skirted twin.tec PCR Plates 96 (Eppendorf) where each well contains 8 μL of RNase-free water containing 0.5 U of protector RNase-inhibitor (Roche). After being deposited into the destination wells, the cells were lysed by osmotic shock. The resulting cell lysate was divided into duplicates, one for scMS and the other for scRNAseq, both kept in –80 °C for short term storage.

### Single cell mass spectrometry

#### Metabolomics profiling of single cell using UPLC-MS

UPLC-MS analysis was performed as described previously on a Vanquish (Thermo Fisher Scientific) system coupled to a Q-Exactive Plus Orbitrap (Thermo Fisher Scientific) mass spectrometer (12). Briefly, chromatographic separation was carried out on a Waters™ ACQUITY UPLC BEH C18 130 Å column (1.7 μm, 1 mm x 50 mm) maintained at 40 °C. The binary mobile phases were 0.1% HCOOH (formic acid) in MilliQ water (aqueous phase) (A) and acetonitrile (ACN) (B). The gradient started with 1% B for 0.5 min, increased linearly to 70% B over 5 min, followed by a wash stage was performed at 99% B for 0.5 min before switching back to 1% B for 1.5 minutes to condition the column for the next injection. Total chromatographic run time was 7 minutes with a flow rate of 0.3 mL min^-1^. The injection volume was 4 μL and the autosampler was kept at 10 °C throughout the analysis.

The Q-Exactive Plus Orbitrap mass spectrometer (Thermo Fisher Scientific) was equipped with a heated electrospray ionization (HESI) source. Acquisition was performed in full scan MS mode (resolution 70,000-FWHM at 200 Da) in positive mode over the mass range *m/z* from 120 to 1,000. The full-scan and data-dependent MS/MS mode (full MS/dd-MS^2^ Top10) was used for QC pooled samples to simultaneously record the spectra of the precursors as well as their MS/MS (fragmentation). In addition, the full MS/dd-MS^2^ mode with inclusion list was also applied for the pooled QC samples to confirm fragments of the selected precursors. The parameters for dd-MS^2^ were set up as follows: resolution 17,500, mass isolation window 0.7 Da, and normalized collision energy (NCE) was set at 3 levels: 15%, 30%, and 45%. Spectrum data format was centroid and all the parameters of the UHPLC-HRMS system were controlled through Xcalibur software version 4.3.73.11 (Thermo Fisher Scientific).

#### LC-MS data processing and analyzing

For targeted analysis, peak area from extracted ion chromatograms (EIC) were integrated and extracted using the Xcalibur Quan Browser version 4.3.73.11 (Thermo Fisher Scientific).

#### Quantification for targeted compounds in single cells

The identification of targeted analytes in the samples was confirmed by comparing retention times to authentic reference standards. Standard solutions were prepared in MeOH at an approximate concentration of 1 mM, with the exact concentration recorded. These solutions were serially diluted down to 0.001 nM and analyzed by UHPLC-MS to determine limit of quantification (LOQ) and calibration range (Table S1). Each calibration point was measured in triplicate, and linear regression curves were calculated using peak areas.

Quantification of analytes in the scMS cell lysate was performed using external calibration curves. Since the lysate was split in half, one for scMS and one for scRNAseq, the analyte amount measured in one portion (for scMS) was doubled to calculate the total amount in a single protoplast. The analyte concentrations in each protoplast were then calculated by dividing the absolute amounts of analytes detected by the estimated volume of the cell, which was calculated assuming the protoplasts are spherical. Protoplast diameters were measured using ImageJ software, based on images obtained from the cell picking robot.

### Single cell RNA sequencing

#### Sequencing library construction and read alignment

Single cell mRNA sequencing libraries were prepared with a half of cell lysate using the SMART-Seq mRNA LP (Takara, 634771) and the customized automation program on Biomek i7 hybrid liquid hander (Beckman Coulter, B87585) according to standard manufacturer’s protocol (PCR 1: 18 cycles and PCR 2: 16 cycles, individually purified). The final library quantification and quality check were performed using Qubit Flex Fluorometer (Thermo Fisher Scientific, Q33327), 4200 TapeStation System (Agilent, G2991BA), and LightCycler 480 II (Roche, 05015278001). The libraries were sequenced on Illumina NovaSeq 6000 (2 × 60 bp) and demultiplexed by bcl2fastq Conversion Software (Illumina) to obtain FASTQ data files.

Index sequences for each fastq file were verified whether they match their corresponding 96-well positions. After confirming qualities of sequencing files via fastqc, adaptors, poly A tails, and low-quality bases were trimmed out of sequencing reads using fastp (28). The cleaned reads were aligned to reference genome (cro_v3) via STAR (v.2.7.10a) and quantified using RSEM (v.1.3.1) (29, 30). The output expected counts were combined into one matrix and subjected to Seurat (v5.0.1) for downstream analyses (31).

#### Processing single-cell transcriptome

Empty wells or duplets were excluded, and cells containing more than 1,000 genes and less than 10,000 genes were kept for downstream analyses. Read counts were log-normalized and 500 variable genes were selected by vst method and scaled for transcriptome-guided dimensional reduction. Genes previously validated by RNA *in situ* hybridization worked as markers for cell type annotation (*NLTP2, CB21, D4H, ISY, G8H*). Cells that have detectable *G8H* or *ISY* were assigned as IPAP cells, and cells that express *D4H* or *DAT* were regarded as idioblasts.

### Comparing transcriptome and metabolome

#### Integration

Before integrating the transcriptomic and metabolomic datasets, empty wells or wells containing doublet of cells were removed based on the images recorded by CellCelector. The estimated concentrations of the targeted metabololites were log_10_-normalized and standardized before appended to Seurat transcriptome object as additional assays. Log-normalized matrices as data and z-scored ones served as scale.data. The presence of marker molecules (secologanin, loganic acid, and serpentine) was used as a marker for metabolite-guided cell type annotation as previously described (12). Concentration of twelve compounds were used as features for metabolite-guided dimensional reduction, and localization of compounds was used for annotating each cluster in metabolite-guided UMAP. Well identity of each cell was utilized to connecting their corresponding UMAP coordinates from transcriptome and metabolome-guided cell clustering.

#### Gene by metabolite correlation

Compound concentration and TPM of genes in each cell was used to calculate Spearman correlation between genes and metabolites. Calculated coefficients of all genes with each specific compound were visualized in scatter plot according to their ascending rank or in heatmap where the coefficients were shown in color scale. Transporters with the top 15 highest and bottom 15 lowest correlation values were connected to corresponding compounds and visualized in the network using Cytoscape (32).

#### Comparing differentially expressed genes between subclusters

Cells accumulating secologanin alone and cells accumulating both secologanin and mauritianin were annotated as each subclusters in epidermis. Differentially expressed genes between two subclusters were extracted using a Wilcoxon Rank Sum test.

### Data, Materials, and Software availability

All data used in this study are available in the article and/or supporting information. Sequencing data have been deposited in NCBI (PRJNA1248169) and are publicly available as of the date of publication. Custom code developed for this work is available in the GitHub repository (https://github.com/moonyoungkang/split_cell_2025).

## Acknowledgements

This work was supported by the Max Planck Society. M.K. acknowledges the National Research Foundation of Korea (RS-2023-00245281) for a grant. We thank Mrs. Eva Rothe and the MPI-CE greenhouse team for growing plants. We thank Ms. Renate Gautsch for her technical support in the NGS Core Facility at MPI-Biochemistry (RRID: SCR_025746). We thank Dr. Mohamed Omar Kamileen, Dr. Gyumin Kang, and Abdullah Sandhu for providing standards used in our analysis. Part of the workflow in Figure 1A was created with BioRender.com.

## Author contributions

M.K., A.H.V., L.C. and S.E.O. designed the study and wrote the paper. M.K., A.H.V., R.K., A.L.C., J.W., S.H. performed the experiments. M.K., A.H.V., A.Y. analyzed the data.

## References

1. P. Berman et al., Parallel evolution of cannabinoid biosynthesis. Nat Plants 9, 817–831 (2023).

2. B. Hong et al., Biosynthesis of strychnine. Nature 607, 617–622 (2022).

3. B. Jiang et al., Characterization and heterologous reconstitution of Taxus biosynthetic enzymes leading to baccatin III. Science 383, 622–629 (2024).

4. C. Y. Kim et al., The chloroalkaloid (-)-acutumine is biosynthesized via a Fe(II)- and 2-oxoglutarate-dependent halogenase in Menispermaceae plants. Nat Commun 11, 1867 (2020).

5. R. S. Nett, W. Lau, E. S. Sattely, Discovery and engineering of colchicine alkaloid biosynthesis. Nature 584, 148–153 (2020).

6. J. Reed et al., Elucidation of the pathway for biosynthesis of saponin adjuvants from the soapbark tree. Science 379, 1252–1264 (2023).

7. I. Efroni, K. D. Birnbaum, The potential of single-cell profiling in plants. Genome Biol 17, 65 (2016).

8. K. H. Ryu, L. Huang, H. M. Kang, J. Schiefelbein, Single-Cell RNA Sequencing Resolves Molecular Relationships Among Individual Plant Cells. Plant Physiol 179, 1444–1456 (2019).

9. M. Kang, Y. Choi, H. Kim, S. G. Kim, Single-cell RNA-sequencing of Nicotiana attenuata corolla cells reveals the biosynthetic pathway of a floral scent. New Phytol 234, 527–544 (2022).

10. C. Li et al., Single-cell multi-omics in the medicinal plant Catharanthus roseus. Nat Chem Biol 19, 1031–1041 (2023).

11. S. Wu, A. L. M. Morotti, J. Yang, E. Wang, E. C. Tatsis, Single-cell RNA sequencing facilitates the elucidation of the complete biosynthesis of the antidepressant hyperforin in St. John’s wort. Mol Plant 17, 1439–1457 (2024).

12. A. H. Vu et al., Quantitative Single-Cell Mass Spectrometry Provides a Highly Resolved Analysis of Natural Product Biosynthesis Partitioning in Plants. J Am Chem Soc 146, 23891–23900 (2024).

13. S. E. O’Connor, J. J. Maresh, Chemistry and biology of monoterpene indole alkaloid biosynthesis. Nat Prod Rep 23, 532–547 (2006).

14. V. Probst et al., Benchmarking full-length transcript single cell mRNA sequencing protocols. BMC Genomics 23, 860 (2022).

15. K. Yamamoto et al., Cell-specific localization of alkaloids in Catharanthus roseus stem tissue measured with Imaging MS and Single-cell MS. Proc Natl Acad Sci U S A 113, 3891–3896 (2016).

16. S. Mahroug, V. Courdavault, M. Thiersault, B. St-Pierre, V. Burlat, Epidermis is a pivotal site of at least four secondary metabolic pathways in Catharanthus roseus aerial organs. Planta 223, 1191–1200 (2006).

17. B. St-Pierre, F. A. Vazquez-Flota, V. De Luca, Multicellular compartmentation of catharanthus roseus alkaloid biosynthesis predicts intercellular translocation of a pathway intermediate. Plant Cell 11, 887–900 (1999).

18. S. Sun et al., Single-cell RNA sequencing provides a high-resolution roadmap for understanding the multicellular compartmentation of specialized metabolism. Nat Plants 9, 179–190 (2023).

19. V. Burlat, A. Oudin, M. Courtois, M. Rideau, B. St-Pierre, Co-expression of three MEP pathway genes and geraniol 10-hydroxylase in internal phloem parenchyma of Catharanthus roseus implicates multicellular translocation of intermediates during the biosynthesis of monoterpene indole alkaloids and isoprenoid-derived primary metabolites. Plant J 38, 131–141 (2004).

20. F. Geu-Flores et al., An alternative route to cyclic terpenes by reductive cyclization in iridoid biosynthesis. Nature 492, 138–142 (2012).

21. C. Li et al., Cell-type-aware regulatory landscapes governing monoterpene indole alkaloid biosynthesis in the medicinal plant Catharanthus roseus. New Phytol 245, 347–362 (2025).

22. K. Yamamoto et al., The complexity of intercellular localisation of alkaloids revealed by single-cell metabolomics. New Phytologist 224, 848–859 (2019).

23. K. Miettinen et al., The seco-iridoid pathway from Catharanthus roseus. Nature communications 5, 3606 (2014).

24. B. Larsen et al., Identification of Iridoid Glucoside Transporters in Catharanthus roseus. Plant Cell Physiol 58, 1507–1518 (2017).

25. F. Yu, V. De Luca, ATP-binding cassette transporter controls leaf surface secretion of anticancer drug components in Catharanthus roseus. Proc Natl Acad Sci U S A 110, 15830–15835 (2013).

26. X. Mao et al., Single-Cell Simultaneous Metabolome and Transcriptome Profiling Revealing Metabolite-Gene Correlation Network. Adv Sci (Weinh) 12, e2411276 (2025).

27. A. Vu, L. Caputi, S. E. O’Connor, Isotopic Labeling Analysis using Single Cell Mass Spectrometry. bioRxiv, 2025.2004. 2010.647921 (2025).

28. S. Chen, Y. Zhou, Y. Chen, J. Gu, fastp: an ultra-fast all-in-one FASTQ preprocessor. Bioinformatics 34, i884–i890 (2018).

29. A. Dobin et al., STAR: ultrafast universal RNA-seq aligner. Bioinformatics 29, 15–21 (2013).

30. B. Li, C. N. Dewey, RSEM: accurate transcript quantification from RNA-Seq data with or without a reference genome. BMC bioinformatics 12, 1–16 (2011).

31. Y. Hao et al., Dictionary learning for integrative, multimodal and scalable single-cell analysis. Nature biotechnology 42, 293–304 (2024).

32. P. Shannon et al., Cytoscape: a software environment for integrated models of biomolecular interaction networks. Genome research 13, 2498–2504 (2003).

