## Supplementary Information for "Single-cell metabolome and RNA-seq multiplexing on single plant cells"

**This PDF file includes:**

Supplementary Scheme 1

Figures S1 to S11

Tables S1 to S3

SI References

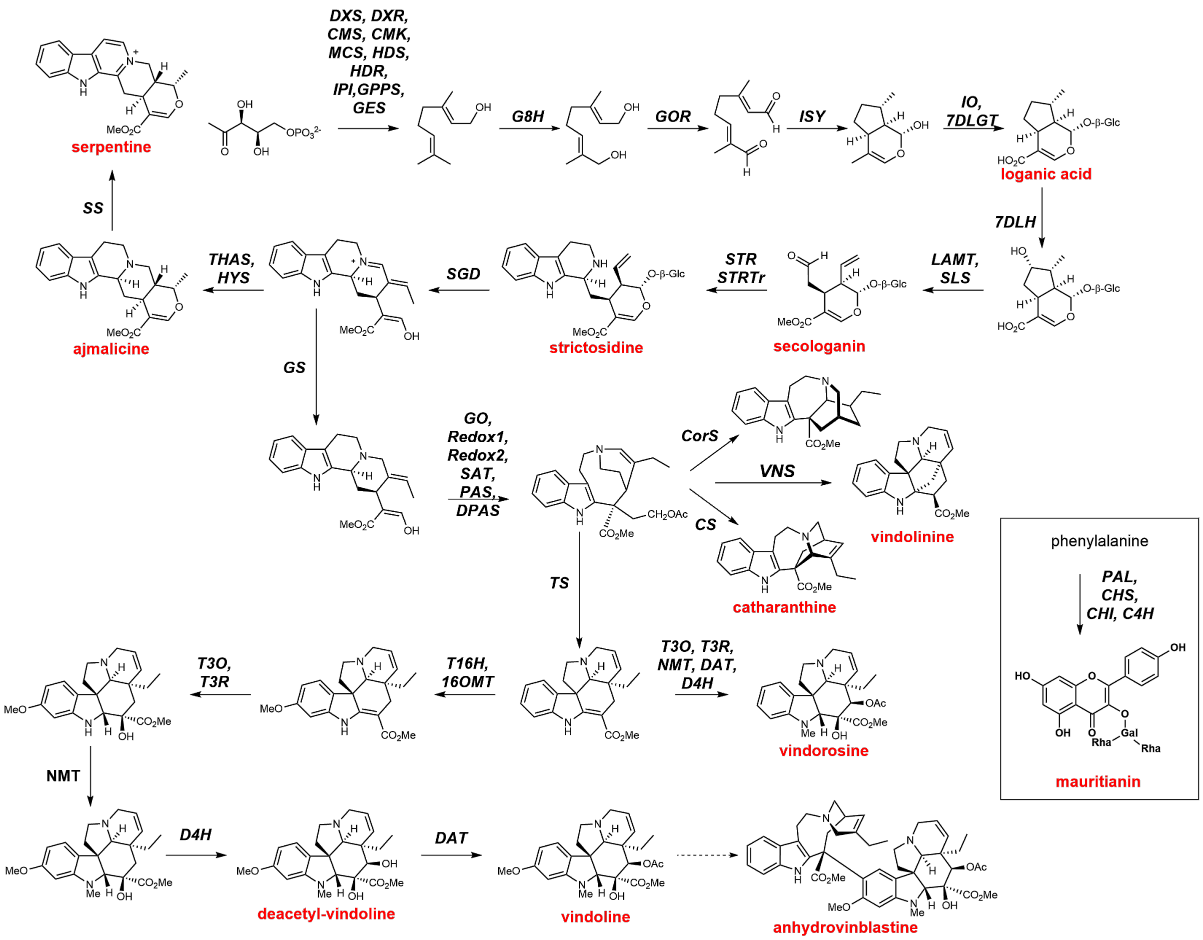


**Supplementary Scheme 1.** A complete biosynthetic pathway to metabolites of *Catharanthus roseus*.

Key metabolites targeted in this study are highlighted in red text. All related biosynthetic genes for these steps are indicated in black, with solid lines. Unconfirmed step is shown as dashed line. The pathway to mauritianin, which is unrelated to other metabolites, is shown inside inset. Gene abbreviations: *DXS*, 1-deoxy-D-xylulose 5-phosphate synthase; *DXR*, 1-deoxy-D-xylulose-5-phosphate reductoisomerase; *CMS*, 4-diphosphocytidyl-methylerythritol 2-phosphate synthase; *CMK*, 4-diphosphocytidyl-2-C-methyl-D-erythritol kinase; *MCS*, 2C-methyl-D-erythritol 2,4-cyclodiphosphate synthase; *HDS*, 1-hydroxy-2-methyl-2-(E)-butenyl-4-diphosphate synthase; *HDR*, 1-hydroxy-2-methyl-butenyl 4-diphosphate reductase; *IPI*, isopentenyl pyrophosphate:dimethylallyl pyrophosphate isomerase; *GPPS*, geranyl pyrophosphate synthase; *GES*, geraniol synthase; *G8H*, geraniol 8-hydroxylase; *GOR*, 8-hydroxygeraniol oxidoreductase; *ISY*, iridoid synthase; *IO*, iridoid oxidase; *7DLGT*, UDP-glucose iridoid glucosyltransferase; *7DLH*, 7-deoxyloganic acid 7-hydroxylase; *LAMT*, loganic acid methyltransferase; *SLS*, secologanin synthase; *STR*, strictosidine synthase; *STRTr*, strictosidine transporter; *SGD*, strictosidine b-glucosidase; *THAS*, tetrahydroalstonine synthase; *HYS*, heteroyohimbine synthase; *SS*, serpentine synthase; *GS*, geissoschizine synthase; *GO*, geissoschizine oxidase; *Redox1*, reductive enzyme 1; *Redox2*, reductive enzyme 2; *SAT*, stemmadenine acetyl transferase; *PAS*, precondylocarine acetate synthase; *DPAS*, dihydroprecondylocarpine synthase; *CorS*, coronaridine synthase; *VNS*, vindolinine synthase; *CS*, catharanthine synthase; *TS*, tabersonine synthase; *T16H*, tabersonine 16-hydroxylase; *16OMT*, 16-hydroxytabersonine O-methyltransferase; *T3O*, tabersonine 3-oxidase; *T3R*, tabersonine 3-reductase; *NMT*, 16-hydroxy-2,3-dihydro-3-hydroxytabersonine N-methyltransferase; *D4H*, desacetoxyvindoline-4-hydroxylase; *DAT*, deacetylvindoline 4-O-acetyltransferase; *PAL*, phenylalanine ammonia lyase; *CHS*, chalcone synthase; *CHI*, chalcone isomerase; *C4H*, cinnamate-4-hydroxylase.

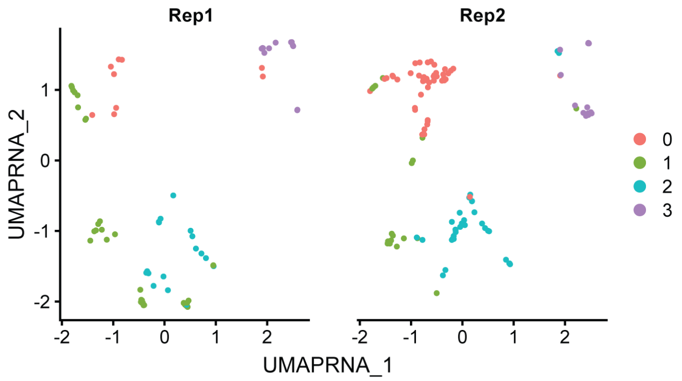


**Fig. S1.** Datasets for single cell RNA sequencing replicates.

After processing single cell RNA sequencing data of cells, their locations were split by original replicate information. The two batches are integrated on UMAP without any significant bias.

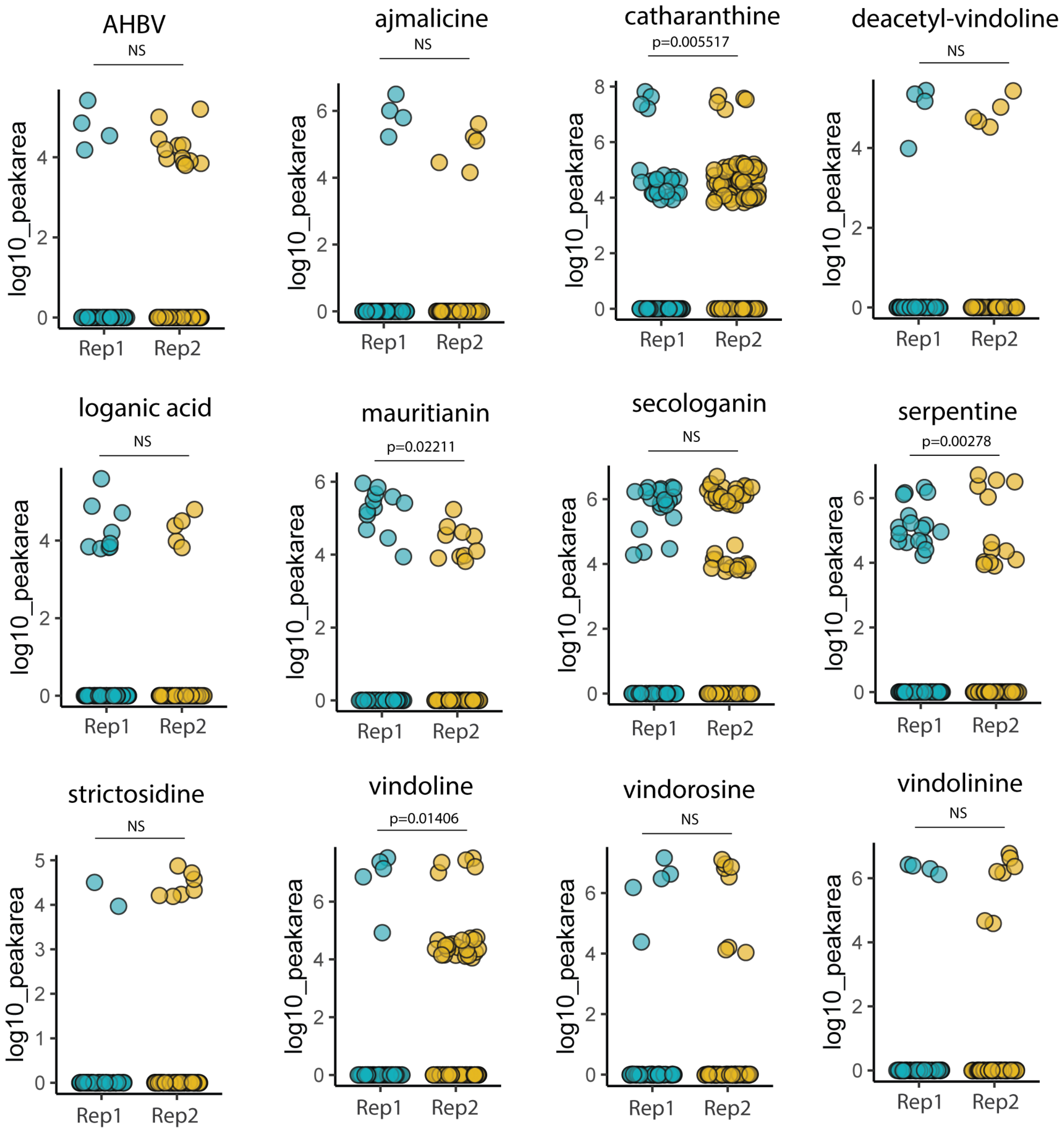


**Fig. S2.** Comparison of the peak areas of targeted analytes from two replicates.

Detected peak areas of targeted analytes were compared between two biological replicates. Most of the analytes did not show a statistical difference between replicates. Numbers of cells that contained vindoline, serpentine, mauritianin, and catharanthine showed differences between replicates, but we assume that this is due to the relatively small number of cells sampled (74 cells in rep1, 119 cells in rep2). Statistical differences between the two replicates were calculated by performing two-sided KS tests (*P* values are annotated; NS, not significant).

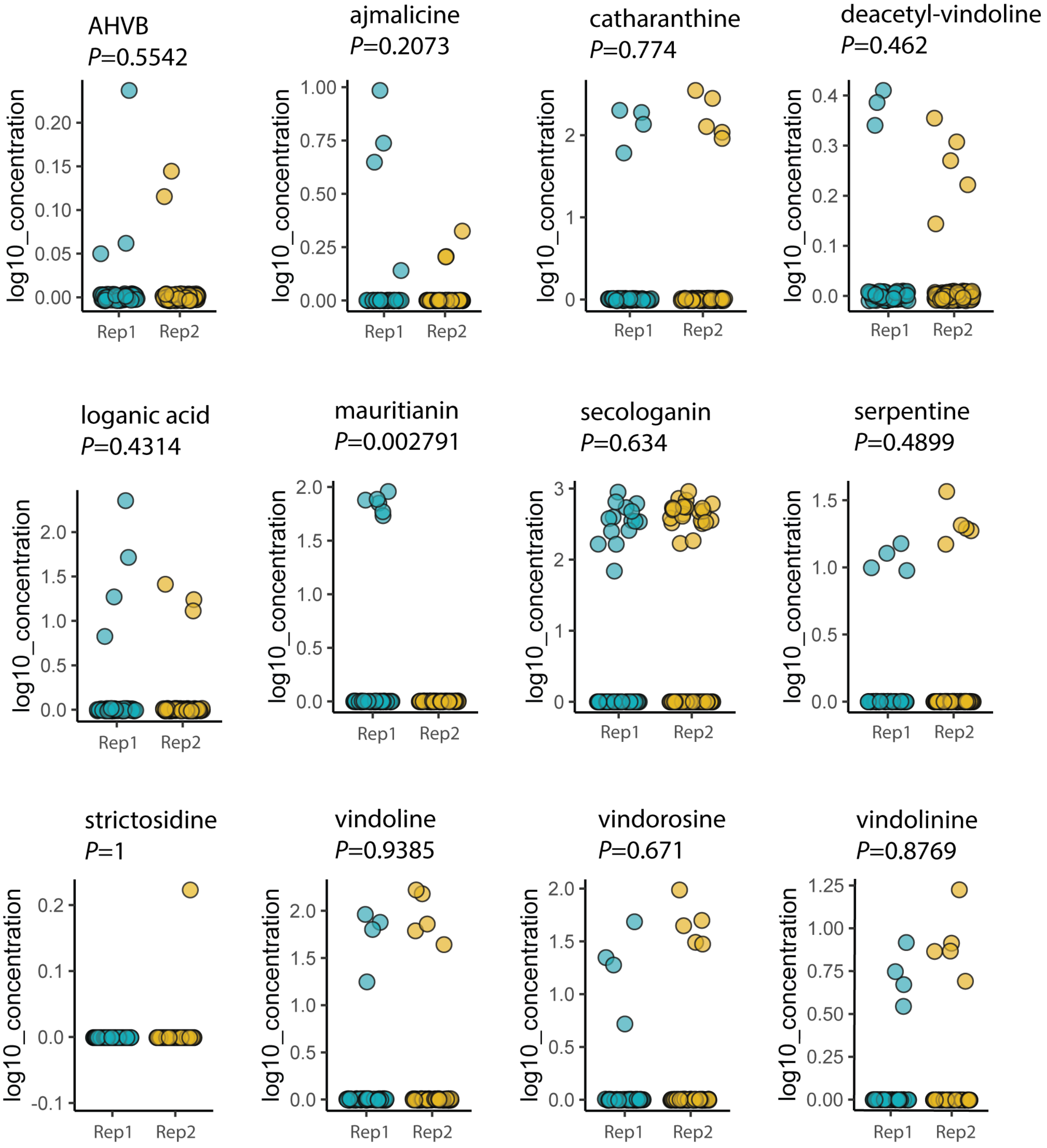


**Fig. S3.** Comparison of the absolute concentration of targeted analytes from two replicates (Concentration)

Calculated intra-cellular concentrations of targeted analytes were compared between two biological replicates. Most of the analytes did not show a statistical difference between replicates. Numbers of cells that contained vindoline, serpentine, mauritianin, and catharanthine showed differences between replicates, but we assume that this is due to the relatively small number of cells sampled (74 cells in rep1, 119 cells in rep2). Statistical differences between the two replicates were calculated by performing two-sided KS tests (*P* values are annotated).

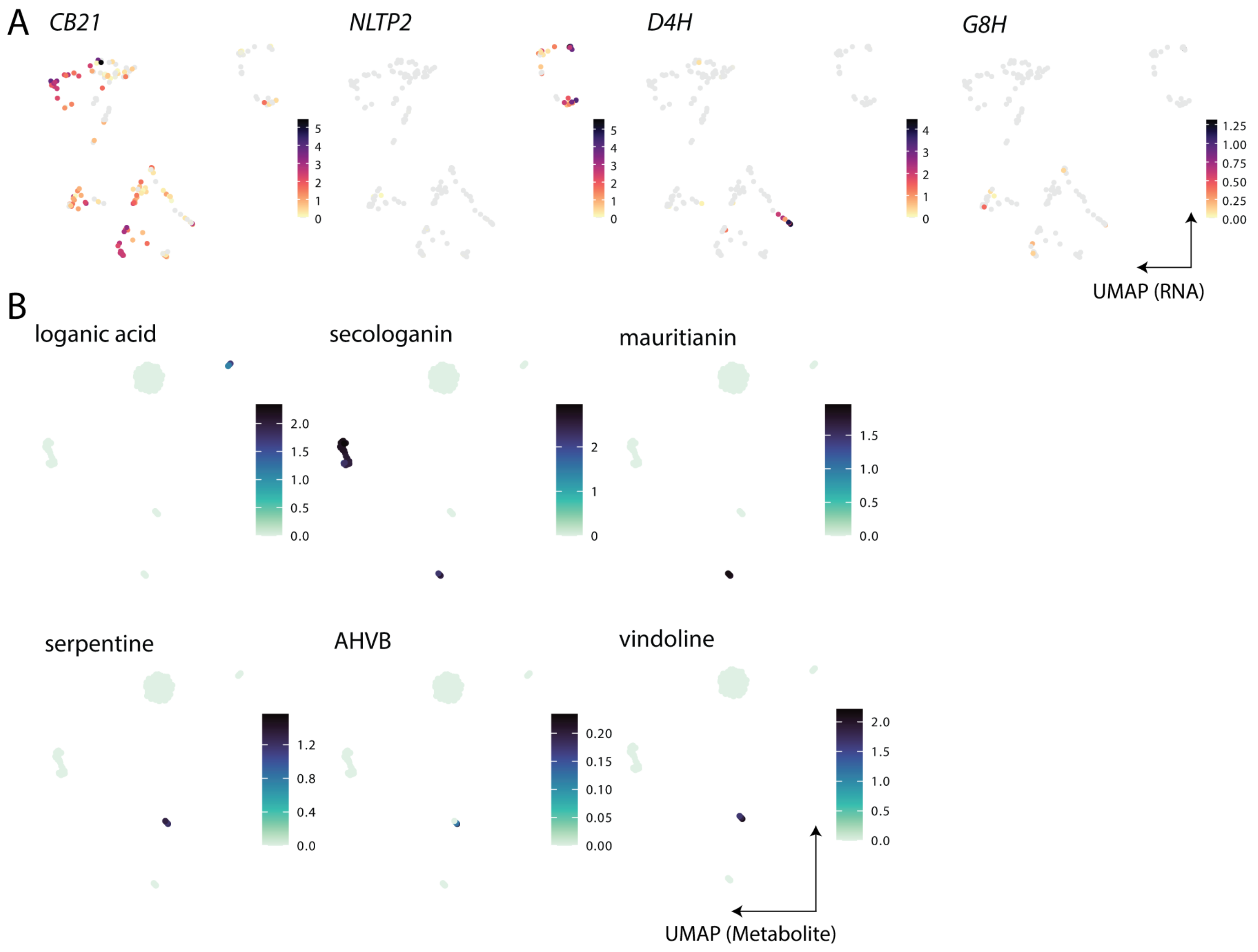


**Fig. S4.** Representative marker genes and analytes that are used for annotating two UMAP plots.

**A.** *CB21* (parenchyma), *NLTP2* (epidermis), *D4H* (idioblast), and *G8H* (IPAP) expressions on RNA UMAP. **B.** loganic acid (IPAP), secologanin and mauritianin (epidermis), serpentine, vindoline, and AHVB (idioblast) accumulation on MET-UMAP.

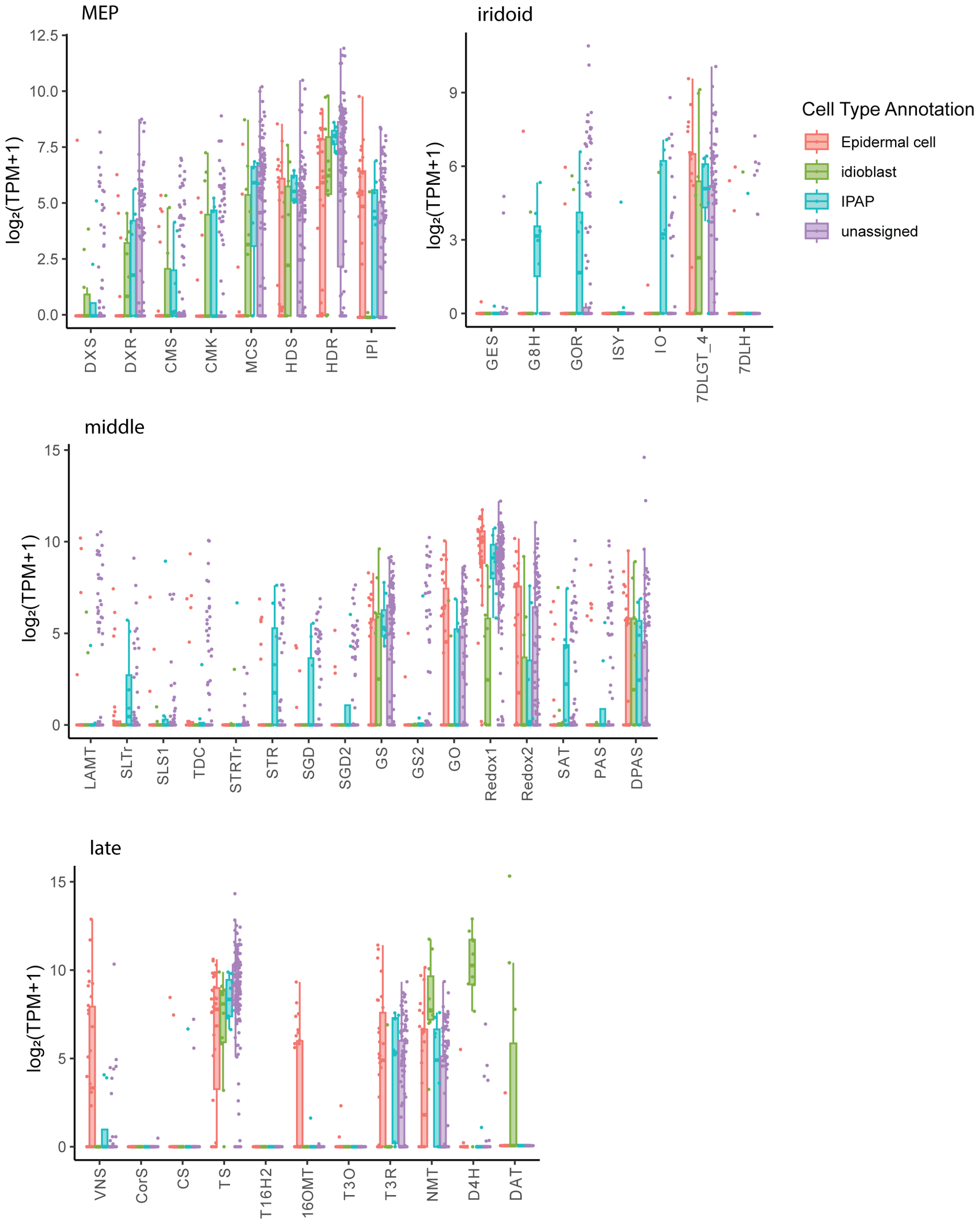


**Fig. S5.** Boxplot showing expression of MIA biosynthetic genes.

Log2-normalized gene expression level is shown as box plot for each gene, grouped and colored by marker gene-guided annotation. The box shows the interquartile range (IQR), and the whiskers correspond to the highest and lowest points within 1.5× IQR.

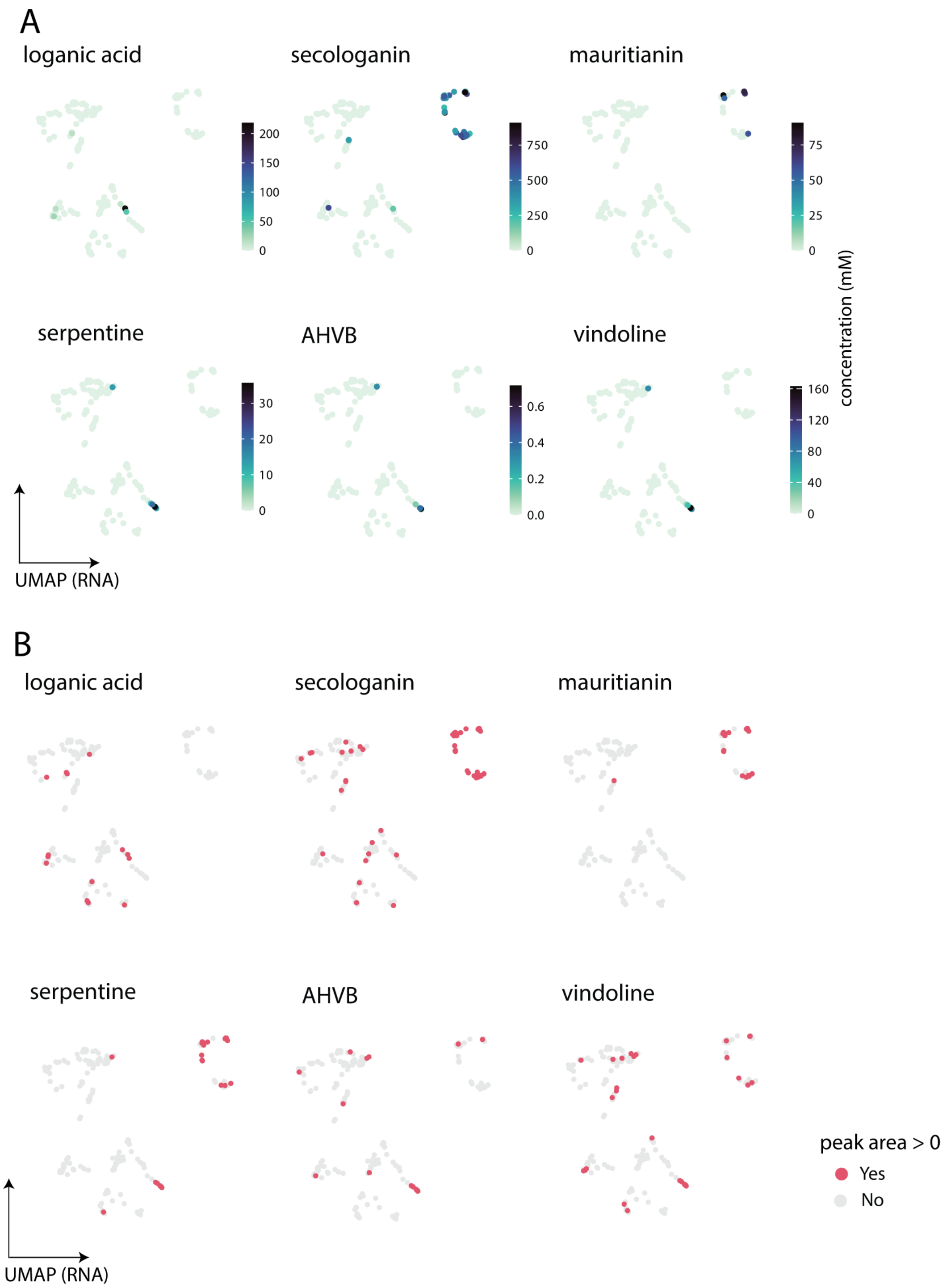


**Fig. S6.** Location of targeted analyte accumulation on RNA-UMAP

**A.** Key analytes are plotted on the RNA-UMAP, showing cell-type specific accumulations. The concentration of each analyte is shown based on color-scale. **B.** Detected peak area are plotted on RNA-UMAP. Secologanin and serpentine show some leaky pattern of accumulation, outside epidermis and idioblast cells, respectively.

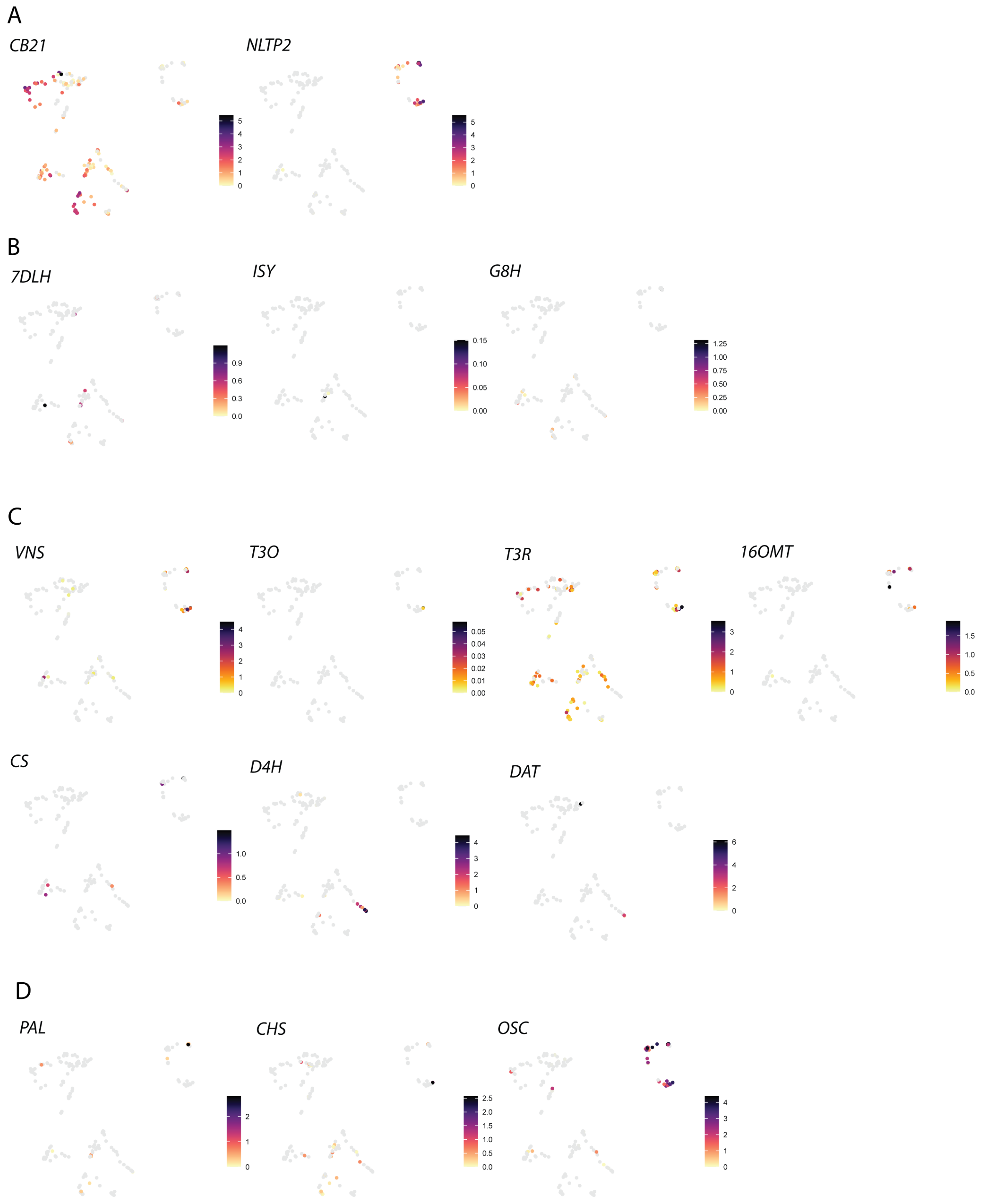


**Fig. S7.** Feature plots of representative genes on RNA-UMAP.

**A.** Parenchyma (*CB21*), epidermis marker (*NLTP2*) genes show exclusive expression pattern in each cell type. **B.** Iridoid biosynthetic show scattered pattern without clear coexistence or clustering. **C.** Late alkaloid biosynthetic genes are mostly localized outside idioblasts, except for *D4H* and *DAT*. **D.** Representative phenylpropanoid genes are localized outside epidermis. *OSC* (Oxidosqualene cyclase) is specifically expressed in epidermis.

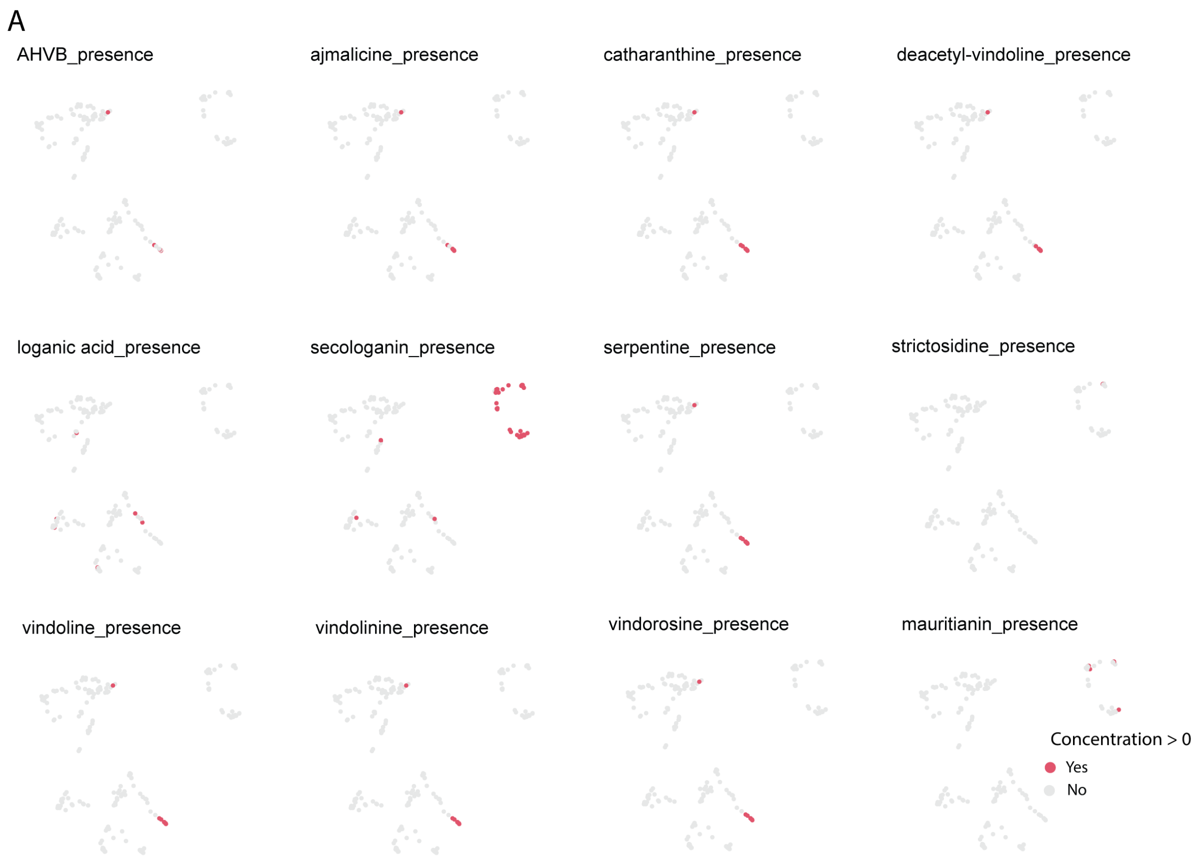


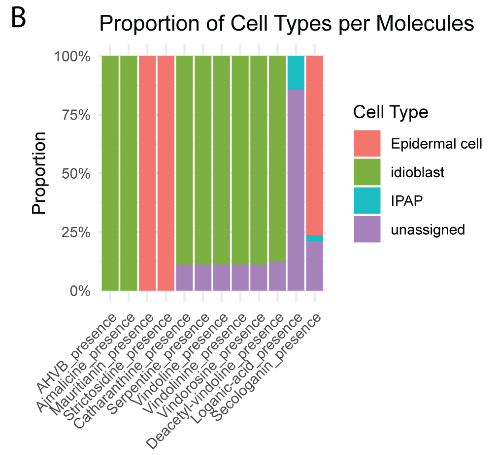


**Fig. S8.** Presence of targeted analytes in each cell type

**A.** Presence of each analyte are plotted as pink dots on RNA-UMAP, when the peak area of each molecule is within quantifiable range. **B.** The stacked bar plot shows proportion of cells that retain each analyte. Cell types are colored based on their RNA-based cell type annotation.

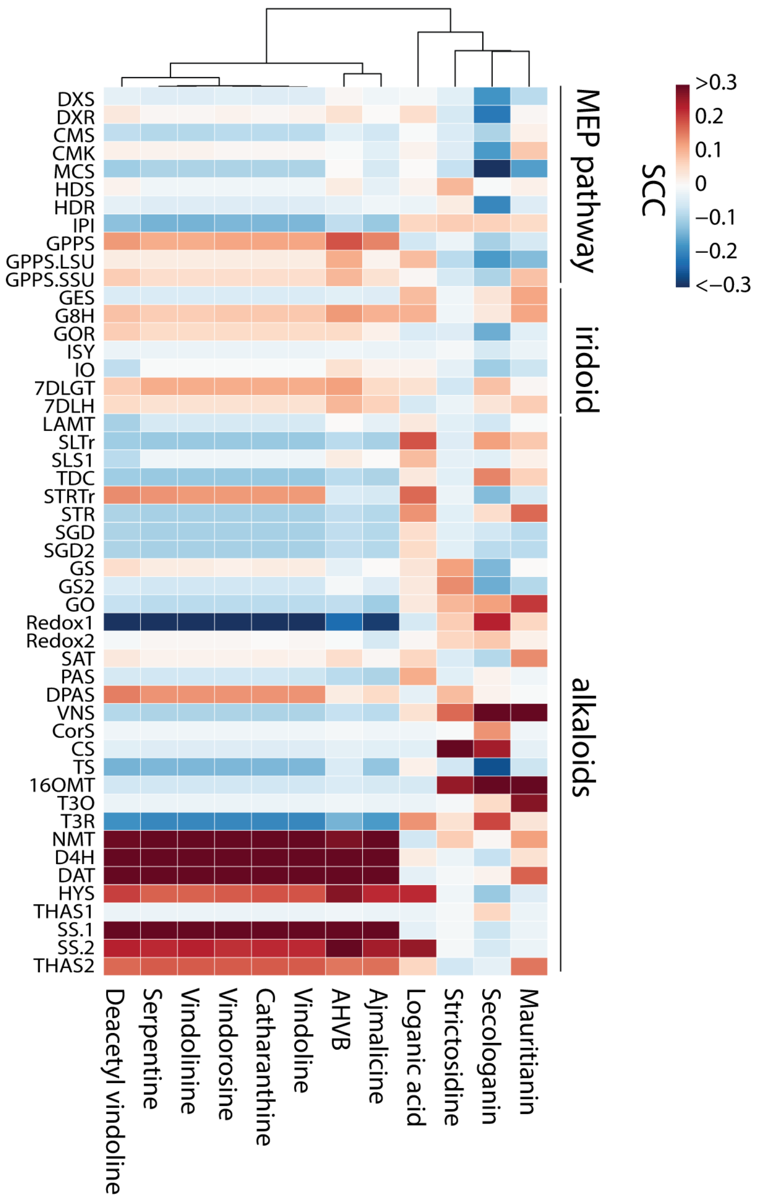


**Fig. S9.** Heatmap showing correlation coefficients of gene and metabolites

Correlation between analytes and known biosynthetic genes are colored based on Spearman correlation coefficients. Late alkaloid genes such as CS, VNS show negative correlation with their products such as Catharanthine Vindoline, suggesting the molecules are actively transported in inter-cellular manner.

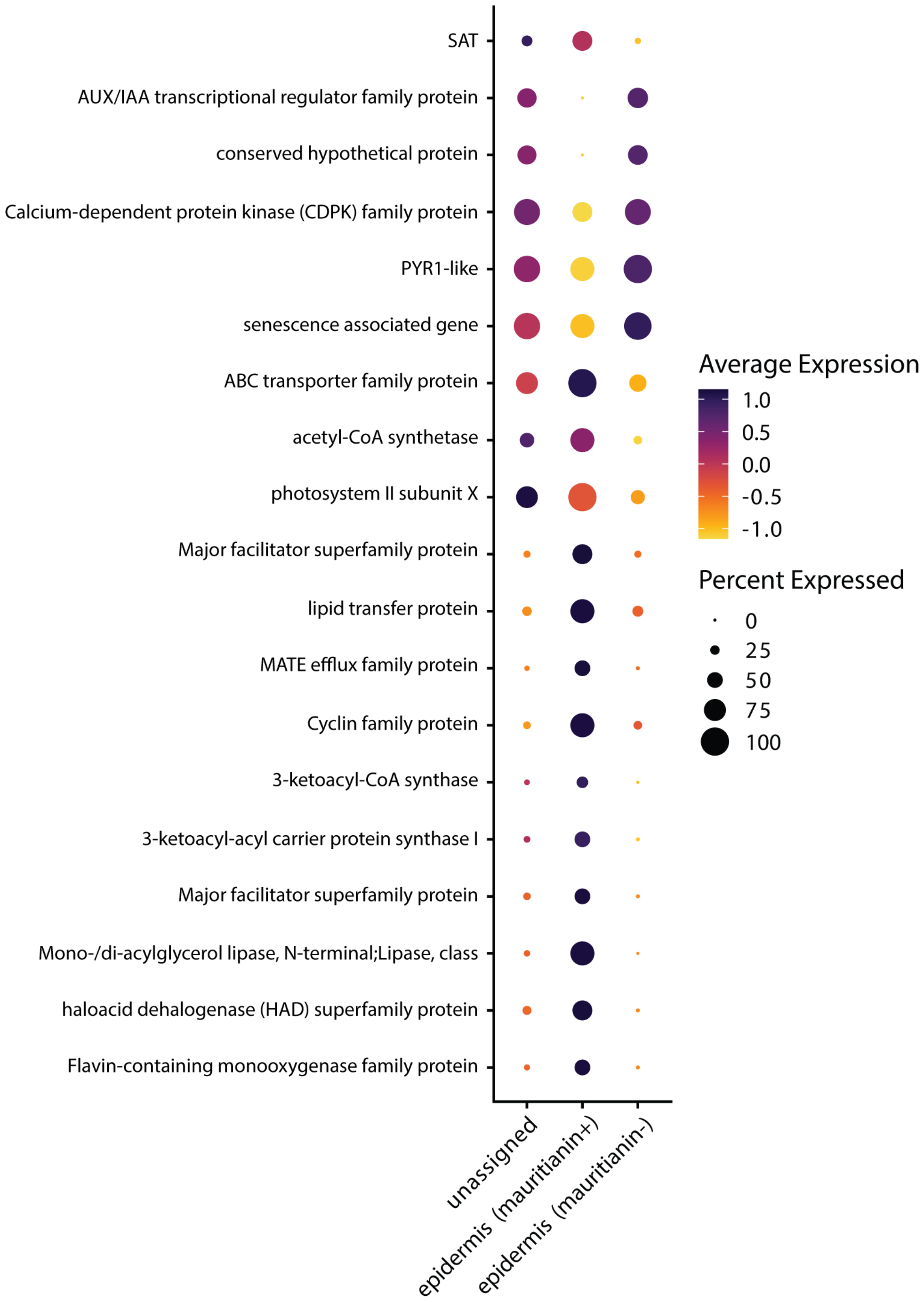


**Fig. S10.** Expression pattern of representative genes that are expressed differentially between two epidermis clusters that are accumulating different analytes**.**

Dotplot shows that epidermal cells that accumulate a quantifiable amount of mauritianin are more likely to be active in lipid metabolism, whereas the other express more senescence or signaling cascade related genes.

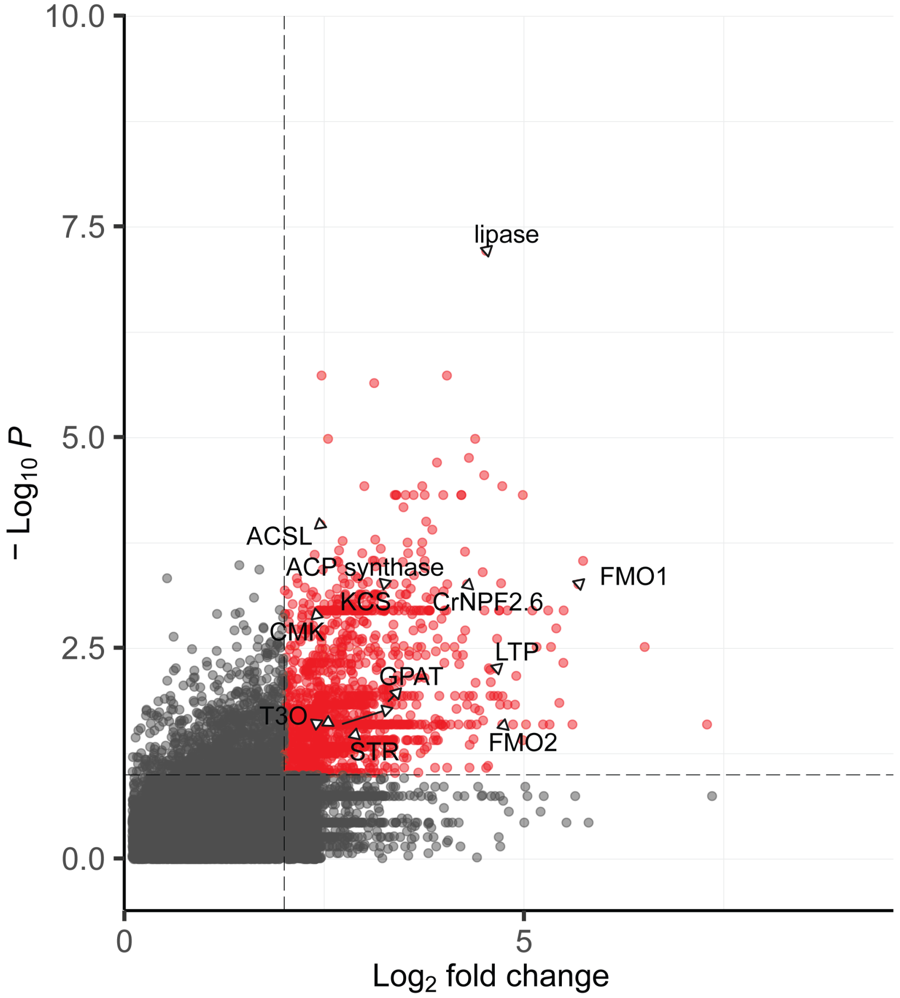


**Fig. S11.** Upregulated genes in epidermal cells that retain quantifiable levels of mauritianin

Scatter plot showing upregulated genes in epidermal cells accumulating a quantifiable amount of mauritianin compared to those without a quantifiable level of mauritianin. Genes with $log_2FC > 2$ and $-log_{10}P$ value are shown in red. ACS synthase, 3-ketoacyl-acyl carrier protein synthase; ACSL, long-chain acyl-CoA synthetase; CMK, 4-(cytidine 5'-phospho)-2-C-methyl-D-erithritol kinase; FMO, Flavin-containing monooxygenase; GPAT, glycerol-3-phosphate acyltransferase; KCS, 3-ketoacyl-CoA synthase; lipase, Mono-/di-acylglycerol lipase; LTP, lipid transfer protein; STR, strictosidine synthase; T3O, tabersonine 3-oxygenase

**Table S1.** Analytical parameters for the external calibration of compounds quantified in this study.

| Compound | Calibration range (nM) | Regression equation[a] | Correlation Coefficient ($R^2$) | LOQ[b] (nM) |
|---|---|---|---|---|
| loganic acid | 2-2000 | y=12248x + 6150 | 0.9999 | 2 |
| secologanin | 10-1000 | y = 16165x + 84261 | 0.9988 | 10 |
| mauritianin | 10-3000 | y = 13958x + 42965 | 0.9998 | 10 |
| strictosidine | 0.1-500 | y = 855571x + 379613 | 0.9999 | 0.1 |
| catharanthine | 0.1-400 | y = 1039954x + 283600 | 0.9997 | 0.1 |
| deacetylvindoline | 0.05-500 | y = 750146x + 1400257 | 0.9985 | 0.05 |
| vindoline | 0.1-500 | y = 1186763x + 1795458 | 0.9991 | 0.1 |
| vindorosine | 0.1-500 | y= 846541x + 597335 | 0.9996 | 0.1 |
| serpentine | 0.5-500 | y = 1196468x + 269956 | 0.9994 | 0.5 |
| vindolinine | 0.05-150 | y=2210449x + 710425 | 0.9993 | 0.05 |
| anhydrovinblastine | 0.02-50 | y = 1976113x + 64437 | 0.9999 | 0.02 |
| ajmalicine | 0.05-200 | y=1013368x+535542 | 0.9993 | 0.05 |

[a]Each point of calibration curve was measured in triplicate.

[b]The LOQ was estimated in the lowest analyte concentration injected that yielded a signal-to-noise (S/N) ratio of ≥10 in three replicates.

**Table S2.** Quantification of targeted analytes by scMS and their occurrence in cells after removing cells with unsuccessful scRNA seq analysis

| Compounds | In this study | | | Vu et al., *JACS*, 2024 (1) | | | Li et al., *Nat. Chem. Bio* (2023)(2) | | |
|---|---|---|---|---|---|---|---|---|---|
| | Mean (mM) | Min-max (mM) | No. cells detected/ No. cells analyzed (%) | Mean (mM) | Min-max (mM) | No. cells detected/ No. cells analyzed (%) | Mean (mM) | Min-max (mM) | No. cells detected/ No. cells analyzed (%) |
| anhydrovinblastine | 0.34 | 0.12 - 0.71 | 5/193 (2.6) | 2.62 | 0.31 - 11.8 | 7/202 (3.5) | 0.3 | 0.01 - 1.4 | 37/627 (5.7) |
| ajmalicine | 2.76 | 0.39 - 8.64 | 7/193 (3.6) | - | - | - | - | - | - |
| catharanthine | 169.35 | 59.27 - 341.05 | 9/193 (4.7) | 99.81 | 72.69 - 126.07 | 5/202 (2.5) | 90.0 | 1.92 - 266.89 | 47/627 (7.5) |
| deacetylvindoline | 1.07 | 0.39 - 1.62 | 8/193 (4.1) | 0.36 | 0.19 - 0.53 | 5/202 (2.5) | - | - | - |
| loganic acid | 49.21 | 5.63 - 218.37 | 7/193 (3.6) | 7.32 | 1.5 - 49.26 | 36/202 (17.8) | - | - | - |
| secologanin | 448.48 | 68.09 - 908.09 | 37/193 (19.2) | 316.92 | 50.96 - 601.07 | 58/202 (28.7) | - | - | - |
| serpentine | 16.50 | 8.47 - 35.66 | 9/193 (4.7) | 4.91 | 2.44 - 7.35 | 5/202 (2.5) | 18.3 | 2.25 - 43.98 | 47/627 (7.5) |
| stemmadenine acetate | - | - | - | - | - | - | - | - | - |
| strictosidine | 0.76 | - | 1/193 (0.5) | - | - | - | - | - | - |
| vinblastine | - | - | - | 0.10 | - | 1/202 (0.5) | 0.2 | 0.08 - 0.52 | 5/627 (0.8) |
| vindoline | 81.58 | 16.85 - 162.99 | 9/193 (4.7) | 53.39 | 32.2 - 80.45 | 5/202 (2.5) | 6.8 | 0.04 - 39.3 | 47/627 (7.5) |
| vindolinine | 6.43 | 2.5 - 15.87 | 9/193 (4.7) | 8.80 | 3.4 - 14.75 | 5/202 (2.5) | - | - | - |
| vindorosine | 37.69 | 4.17 - 96.3 | 9/193 (4.7) | 38.27 | 24.96 - 56.32 | 5/202 (2.5) | - | - | - |
| mauritianin | 69.19 | 52.71 - 90.68 | 5/193 (2.6) | 21.65 | 4.92 - 47.73 | 16/202 (7.9) | - | - | - |
| coronaridine | - | - | - | 0.62 | 0.17 - 2.19 | 5/202 (2.5) | - | - | - |

**Table S3.** Number of cells that were annotated as each cell type

Cell types based on RNA (UMAP plot of Figure 2A) were annotated based on known marker gene expression (*NLTP2* for epidermis, *G8H* or *ISY* for IPAP, and *D4H* or *DAT* for idioblast). MET based annotation was done as previously described in A. H. Vu *et al*. 2024, where secologanin served as marker for epidermis, serpentine did for idioblast, and loganic acid did for IPAP.

| | **RNA based annotation** | | **MET based annotation** | |
|---|---|---|---|---|
| **Cell type** | **Rep 1** | **Rep 2** | **Rep 1** | **Rep 2** |
| epidermis | 14 | 16 | 15 | 23 |
| idioblast | 6 | 4 | 4 | 5 |
| IPAP | 4 | 4 | 4 | 3 |
| parenchyma/unassigned | 50 | 95 | 51 | 88 |

**SI References**


1. A. H. Vu *et al.*, Quantitative Single-Cell Mass Spectrometry Provides a Highly Resolved Analysis of Natural Product Biosynthesis Partitioning in Plants. *J Am Chem Soc* **146**, 23891-23900 (2024).
2. C. Li *et al.*, Single-cell multi-omics in the medicinal plant Catharanthus roseus. *Nat Chem Biol* **19**, 1031-1041 (2023).
